# Dissecting the oncogenic mechanisms of POT1 cancer mutations through deep scanning mutagenesis

**DOI:** 10.1101/2024.08.19.608636

**Authors:** Annika Martin, Johannes Schabort, Rebecca Bartke-Croughan, Stella Tran, Atul Preetham, Robert Lu, Richard Ho, Jianpu Gao, Shirin Jenkins, John Boyle, George E. Ghanim, Milind Jagota, Yun S. Song, Hanqin Li, Dirk Hockemeyer

## Abstract

Mutations in the shelterin protein POT1 are associated with diverse cancers, but their role in cancer progression remains unclear. To resolve this, we performed deep scanning mutagenesis in POT1 locally haploid human stem cells to assess the impact of POT1 variants on cellular viability and cancer-associated telomeric phenotypes. Though POT1 is essential, frame-shift mutants are rescued by chemical ATR inhibition, indicating that POT1 is not required for telomere replication or lagging strand synthesis. In contrast, a substantial fraction of clinically-validated pathogenic mutations support normal cellular proliferation, but still drive ATR-dependent telomeric DNA damage signaling and ATR-independent telomere elongation. Moreover, this class of cancer-associated POT1 variants elongates telomeres more rapidly than POT1 frame-shifts, indicating they actively drive oncogenesis and are not simple loss-of-function mutations.

## Main Text

Mutations in POT1 have been identified in multiple cancer types and are estimated to have an overall prevalence of up to 3% (*1–4*). Cancer-associated POT1 (caPOT1) mutations are missense, splicing, or nonsense mutations which can be found throughout the POT1 gene and are almost exclusively heterozygous. While a small subset of these POT1 mutations have been linked to familial cancer predisposition syndromes (*5–13*), the vast majority, over 900 mutations on ClinVar, are found in sporadic cancers and have been annotated as variants of uncertain significance (VUS). Therefore, a key challenge in understanding the role of POT1 in carcinogenesis is delineating which mutations are pathogenic and by which mechanism these mutations drive the disease. This challenge is complex, especially since POT1 plays key roles in multiple telomeric processes: POT1 protects the telomeric overhang from being detected by the ATR kinase (*14*), controls telomere overhang generation and length (*15*, *16*), limits access of telomerase to the telomere (*17*, *18*), and has been implicated in mediating telomere replication (*19*, *20*).

Despite the number of identified variants, relatively few have been molecularly characterized. Of these, several fall within the first two oligonucleotide-binding (OB) folds of POT1, which are required for POT1 binding to the single-stranded DNA (ssDNA) telomeric overhang (*18*, *21*, *22*). Disrupting POT1 ssDNA binding by deletion of the first OB fold or point mutations at key interacting residues results in telomere elongation (*11*, *23*, *24*), presumably by increasing the accessibility of the overhang to telomerase activity. Other mutations in the C-terminal OB fold of POT1 disrupt the POT1-TPP1 interaction (*25*) which is essential for proper localization of POT1 to telomeres (*26*). These mutations can similarly drive telomere elongation (*27*). Therefore, caPOT1 mutations may drive cancer by forestalling telomere-shortening induced replicative senescence and allowing for further acquisition of driver mutations. Beyond overhang sequestration, POT1 also negatively regulates telomere length via POT1-TPP1 recruitment of the CTC1-STN1-TEN1 (CST) complex (*20*, *28*), which terminates telomerase activity (*29*). However, CST is required for successful fill-in synthesis and telomere maintenance (*30*), and POT1 mutations have also been identified in telomere shortening disorders (*19*, *31*), indicating that POT1’s role in telomere length regulation is complex and multifaceted, further challenging the characterization of POT1 VUS in cancer development.

A central outstanding question about telomere length control by POT1 and telomerase action at telomeres is whether it can be separated from DNA damage signaling at telomeres. In some POT1 cancer models, telomere elongation is accompanied by deprotection of telomere ends and aberrant activation of DNA damage response (DDR) pathways at the telomere, indicating that caPOT1 mutations may also drive cancer by triggering genomic instability (*16*, *24*, *32*, *33*). In cell culture assays, depletion of POT1 triggers activation of ATR kinase and the accumulation of fragile telomeres, branched telomeric DNA structures, and extrachromosomal telomeric DNA indicative of failed telomere maintenance (*16*, *34*, *35*). However, this phenotype is not easily uncoupled from telomere length regulation. Interestingly, even in healthy cells, POT1’s protective function seems to be lost for a short period at the end of S-phase as both ATR and ATM signaling can be detected at chromosome ends (*36–38*). As initially established in yeast (*39–41*), ATR and ATM signaling may also control telomere length in human cells, as knockdown of either pathway reduces recruitment of telomerase (*38*).

The characterization of caPOT1 mutations is further challenged by the broad distribution of caPOT1 mutations across the gene body rather than precise clustering into functionally defined regions. Thus, it remains unclear by which mechanism, or combination of mechanisms, POT1 variants drive cancer progression. The occurrence of early truncating mutations and variants which disrupt the starting methionine indicate that the levels of POT1 expression within the cell play a role in carcinogenesis and that POT1 may function as a haploinsufficient tumor suppressor gene (*33*). Under this hypothesis, that POT1 mutations drive cancer through loss-of-function alleles, we have performed a detailed functional analysis of POT1 through deep scanning mutagenesis to evaluate the effects of single amino acid deletions, alanine substitutions, and more than 600 POT1 VUS. As caPOT1 mutations can be inherited or occur early in tumor progression and are believed to drive disease progression prior to DNA damage checkpoint inactivation (*33*, *42*), the functional annotation of POT1 variants requires a primary cell system which is sensitive to the activation of ATR-mediated DDR and retains an intact telomere maintenance mechanism. Therefore, to enable direct genotype-to-phenotype characterization in bulk, we utilized a recently-described method of generating locally-haploid (loHAP) hESCs to perform our mutagenesis screen and follow up experiments (*43*). Isolation and characterization of key dissociation-of-function mutations and comparison to frame-shift clones demonstrate that cancer associated POT1 mutations are not mere loss-of-function mutations but instead actively promote telomere elongation.

### Generating a deep mutational scanning map for POT1 in human pluripotent stem cells

POT1 loHAP cells were generated from diploid hESCs via a 1.6Mb excision of one POT1 allele including upstream regulatory regions using CRISPR/Cas9 (Fig. 1A). Successful loHAP generation was confirmed via junction sequencing and clonality determined by next generation sequencing (NGS) of a single nucleotide polymorphism (SNP) on the remaining allele (fig. S1A). This enabled us to mutagenize the remaining POT1 allele in POT1 loHAPs and evaluate the effect of these secondary mutations over time by quantifying changes in allele frequency as previously described for the BRCA2 locus (*43*) (Fig. 1A). As expected for an essential gene, frame-shift mutations in POT1 loHAPs were strongly depleted within three weeks of editing compared to the diploid parental, while the allele frequency of unedited and synonymous mutations proportionally increased (Fig. 1B and fig. S1, B and C).

**Fig. 1.**
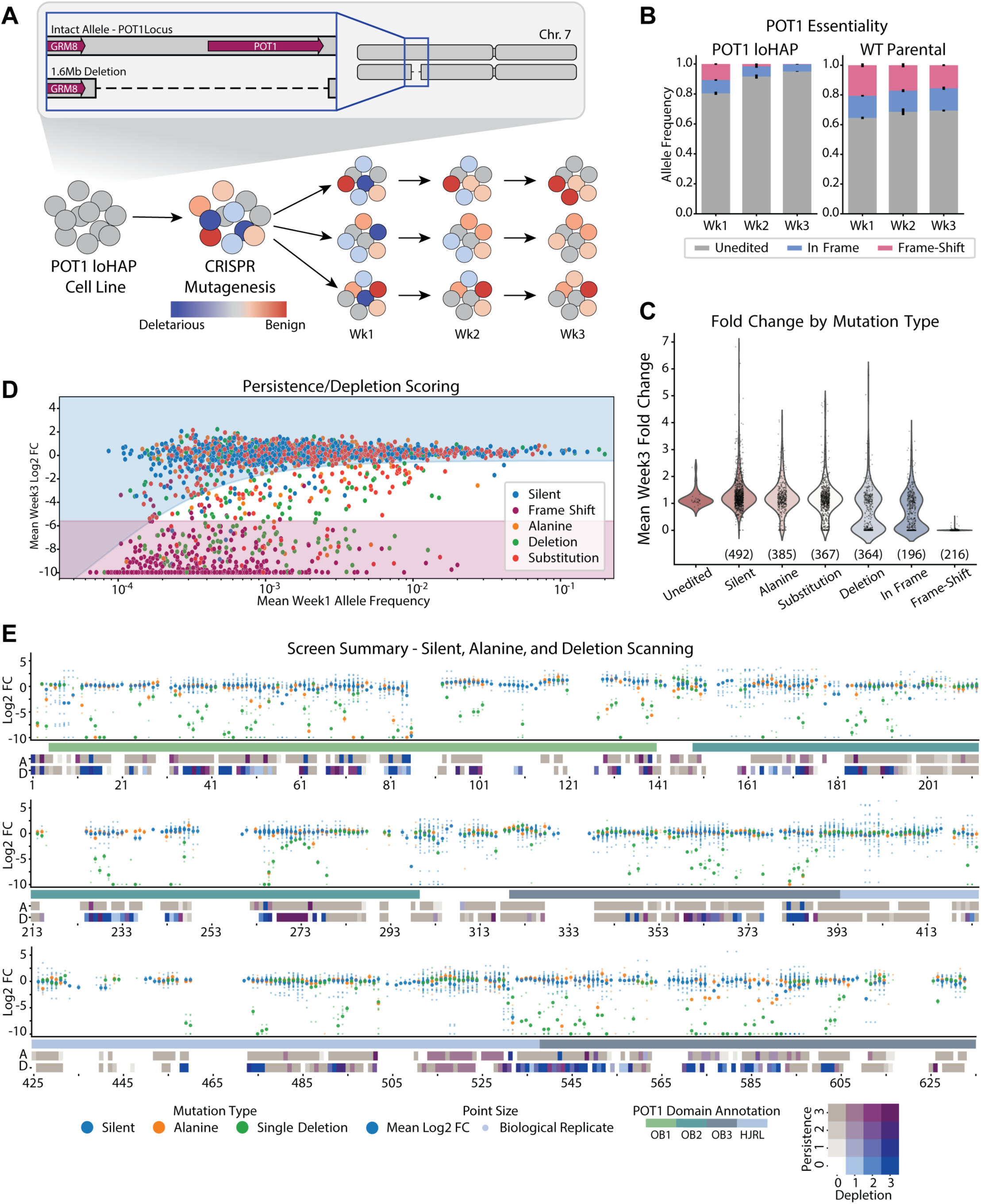
Deep scanning mutagenesis of POT1 reveals essential amino acids in POT1. (**A)** Schematic of POT1 loHAP generation and screen timeline. HDR-mediated CRISPR/Cas9 mutagenesis of the remaining POT1 allele results in a spectrum of mutations with a direct genotype to phenotype correlation. Over three weeks of culture, the most deleterious alleles deplete from the population. (**B**) Relative quantification of CRISPR/Cas9-mediated insertion and deletion alleles over three weeks in POT1 loHAP and diploid wild-type parental cell lines; three biological replicates, with error bars representing the standard error of the mean. (**C**) Mean fold-change in allele frequency across three biological replicates per allele between week 3 and week 1, stratified by mutation type: unedited, silent mutations, alanine substitutions, other substitutions including clinical variants, precise single-codon deletions, other in-frame NHEJ events, and frame-shift NHEJ events. (**D**) Comparison of allele frequency and week 3 log2 fold-change for indicated mutation types. The shaded curves show the 95th percentile for log2 fold-change for silent mutations (blue) and frame-shift mutations (magenta), which were used as boundaries for assigning persistence and depletion scores. (**E**) Summary of the week 3 log2 fold-change for all alanine, silent, and single-codon deletions, with the large dots indicating mean values at a given amino acid position and smaller dots showing individual biological replicates. Below the dot plot is a domain map, showing the location of the HJRL and OB folds. Below the domain map, persistence/depletion scores (see Materials and Methods section Data Analysis - Allele Classification) for alanine substitutions (A) and single-codon deletions (D) are shown for each amino acid position. White (0,0) indicates that the allele was not generated or that the allele could not be classified using these metrics.

To precisely evaluate caPOT1 mutations, we optimized protocols for homology-directed repair (HDR) mediated introduction of designer mutations in POT1 using single-stranded oligonucleotide pools as repair templates (fig. S1, D and E). Using this approach, we successfully generated mutations at 535 of 634 amino acid positions within POT1 (84.4%), including alanine substitutions at 385 unique positions (63.0%; POT1 naturally contains 23 alanine residues) and single-codon deletions at 364 unique positions (57.4%), with synonymous mutations at 492 unique positions (79.8%; 18 residues have no possible synonymous mutations) to act as controls. Allele depletion was evaluated by the fold-change of three independent mutant pools from week one to week three (Fig. 1C and table S1) compared to the behavior of synonymous and frame-shift alleles. To aid in visualization, alleles were also assigned a persistence/depletion score based on the behavior of individual biological replicates (Fig. 1D and fig. S1F). In general, alanine substitutions were more tolerated than single-codon deletions, though among all mutations surveyed, allele depletion was strongest at the N-terminal portion of OB1 (aa1-85) and the C-terminal portion of OB3 (aa538-634) (Fig. 1 E and table S1).

### Deep scanning mutagenesis of POT1 reveals essential amino acids in POT1 stability and function

Alanine and deletion scanning, coupled with existing structural models for POT1 and its interacting partners, TPP1 and telomeric DNA, allowed us to generate a detailed map of residues essential to POT1 function and/or structural stability (*22*, *44–46*). Deletions which result in statistically significant depletion relative to synonymous mutations tend to cluster into contiguous stretches that largely correspond to secondary structural elements (Fig. 2A); Overall, 54% of single deletions within alpha helices and 82% of deletions within beta strands deplete, compared to 34% of deletions in unstructured regions (based on JPRED4 structural classification (*47*) using a minimum confidence threshold of 3). In contrast, depleted alanine substitutions were less clustered and more dispersed along the peptide chain (fig. S2A). Notably, our screen demonstrates that C382 and C385 are essential, as both alanine substitution and deletion resulted in strong depletion (Fig. 2B). These residues, with C503 and C506, are proposed to coordinate a Zinc ion (*22*, *44*, *46*). Therefore, mutations at these residues which disrupt Zinc coordination may result in misfolding of POT1’s Holliday junction resolvase-like domain (HJRL) and OB3. Likewise, alanine substitutions of large hydrophobic amino acids in the core of OB1 and OB2 were strongly selected against, suggesting that alanine substitution abrogates essential hydrophobic interactions within the OB cores, resulting in compromised protein stability (Fig. 2C).

**Fig. 2.**
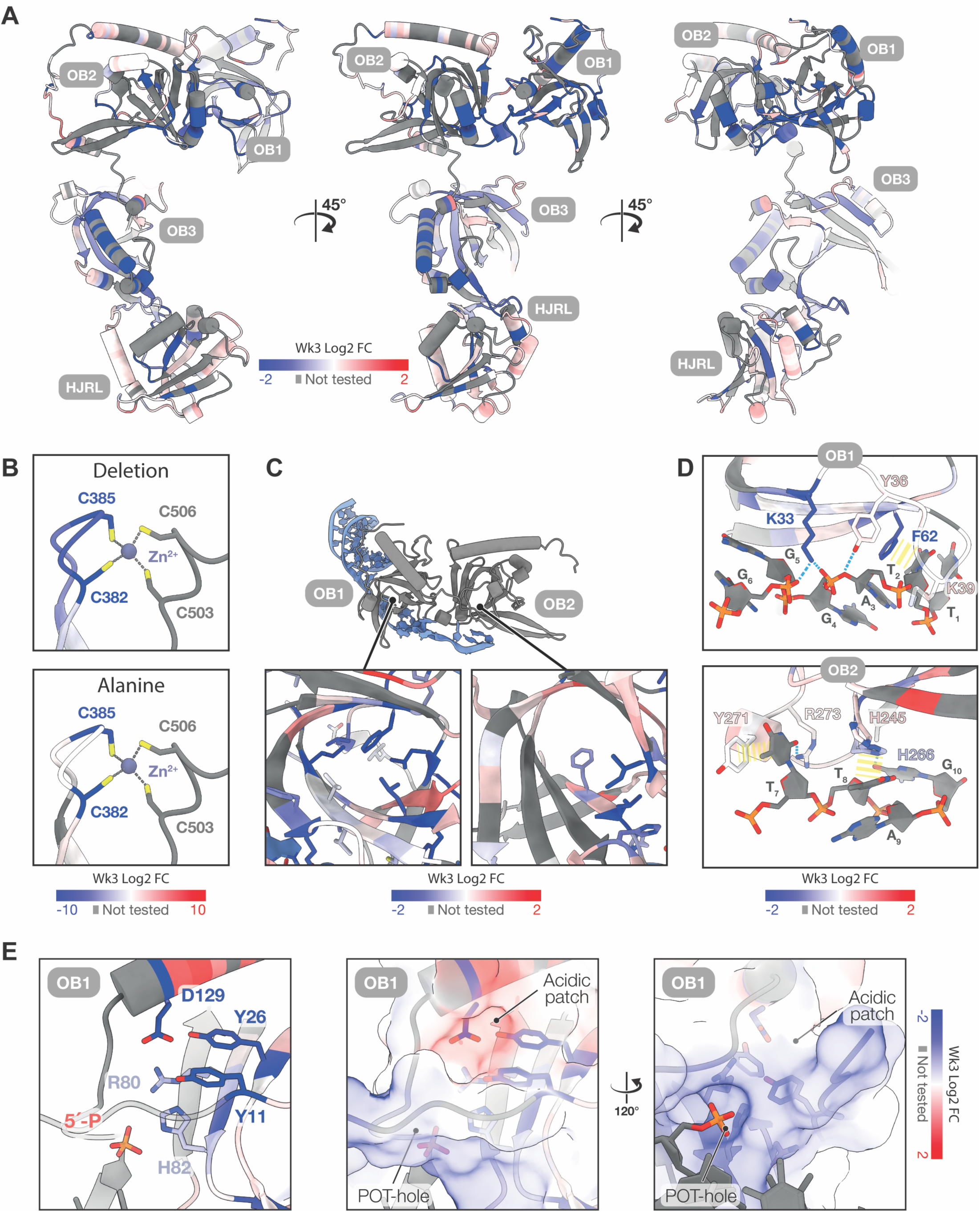
Essential POT1 residues map to structurally defined regions of POT1. (**A**) POT1 AlphaFold2 (*60*) prediction colored by the week 3 log2 fold-change in allele frequency of single-codon deletions at the respective amino acid position. (**B**) Close view of zinc-coordination by Cysteines 382, 385, 503 and 506, colored by the week 3 log2 fold-change in allele frequency of single-codon deletions (top) or alanine substitutions (bottom). Model derived from PDB: 5UN7(*44*). (**C**) POT1 (gray) interaction with telomeric DNA (light blue). Inset windows show hydrophobic residues at the center of OB1 (left) and OB2 (right) colored by log2 fold-change in allele frequency of alanine substitutions. Model derived from PDB: 8SH1 (*45*). (**D**) Closer view of POT1-ssDNA interactions in panel C with highlighted POT1 residues colored by log2 fold-change in allele frequency of alanine substitutions. Model derived from PDB: 1XJV (*22*). (**E**) Closer view of the POT-hole (left) and adjacent acidic patch (center) in OB1, with highlighted POT1 residues colored by log2 fold-change in allele frequency of alanine substitutions. Surface is colored by charge, as per convention where blue and red colors represent the positive and negative charge scores, respectively. The view of the acidic patch (center) is rotated to view the POT-hole (right). Model derived from PDB: 8SH1(*45*).

These findings stand in contrast to mutations of POT1 residues which directly interact with single-stranded telomeric DNA (Fig. 2D) (*22*, *45*). Several alanine substitutions at ssDNA-interacting residues of OB1 do show robust depletion, including F62 which directly stacks against base T1 of the telomeric repeat sequence, and K33 which interacts with the phosphate backbone of the ssDNA (*22*). However, Y36 substitution which also contacts the ssDNA backbone does not deplete. Moreover, alanine substitutions of key amino acids in OB2 do not significantly impair cell viability over the course of three weeks, including H245, H266, Y271, and R273 which make numerous interactions with single-stranded telomeric DNA (*22*). Mutations in the recently described POT-hole (*45*) also persist (Fig. 2E and fig. S2, B and C), including residues R80, R82, and R83 which form a positively charged pocket contacting the 5’ phosphate at the ssDNA-dsDNA junction (*48*). While previous *in vitro* experiments show that alanine substitutions at these positions dramatically reduce POT1 binding to the ssDNA-dsDNA junction of a model telomeric substrate (*45*), this interaction does not appear to be essential in our stem cell system. Interestingly, a solvent-exposed acidic patch adjacent to the POT-hole does show strong depletion, though its potential function is yet undefined (Fig. 2E).

### Many caPOT1 mutations are not simple loss-of-function mutations and fully support cellular proliferation

The critical finding of our alanine scanning, that substitution of key DNA-interacting residues does not necessarily reduce cellular viability, is surprising since mutant POT1’s impaired ability to bind ssDNA, driving subsequent telomere deprotection, has been interpreted as a key feature by which caPOT1 mutations drive carcinogenesis. Thus, to better understand the effect of caPOT1 mutations, we evaluated an additional 624 missense mutations at 367 unique positions which are VUS listed on the ClinVar database or are associated with POT1 cancer through familial inheritance pattern or molecular characterization (*1*, *2*, *5–12*). Of interest, not all substitution mutations at the same amino acid position showed equivalent depletion, and depletion of alanine substitutions was not necessarily predictive of VUS behavior (Fig. 3A). CaPOT1 mutations frequently showed no effect on cellular proliferation or only a mild survival phenotype that fell between synonymous mutations and frame-shift mutations (Fig. 1D), suggesting that many of the variants of unknown significance in POT1 retained the ability to protect telomeres.

**Fig. 3.**
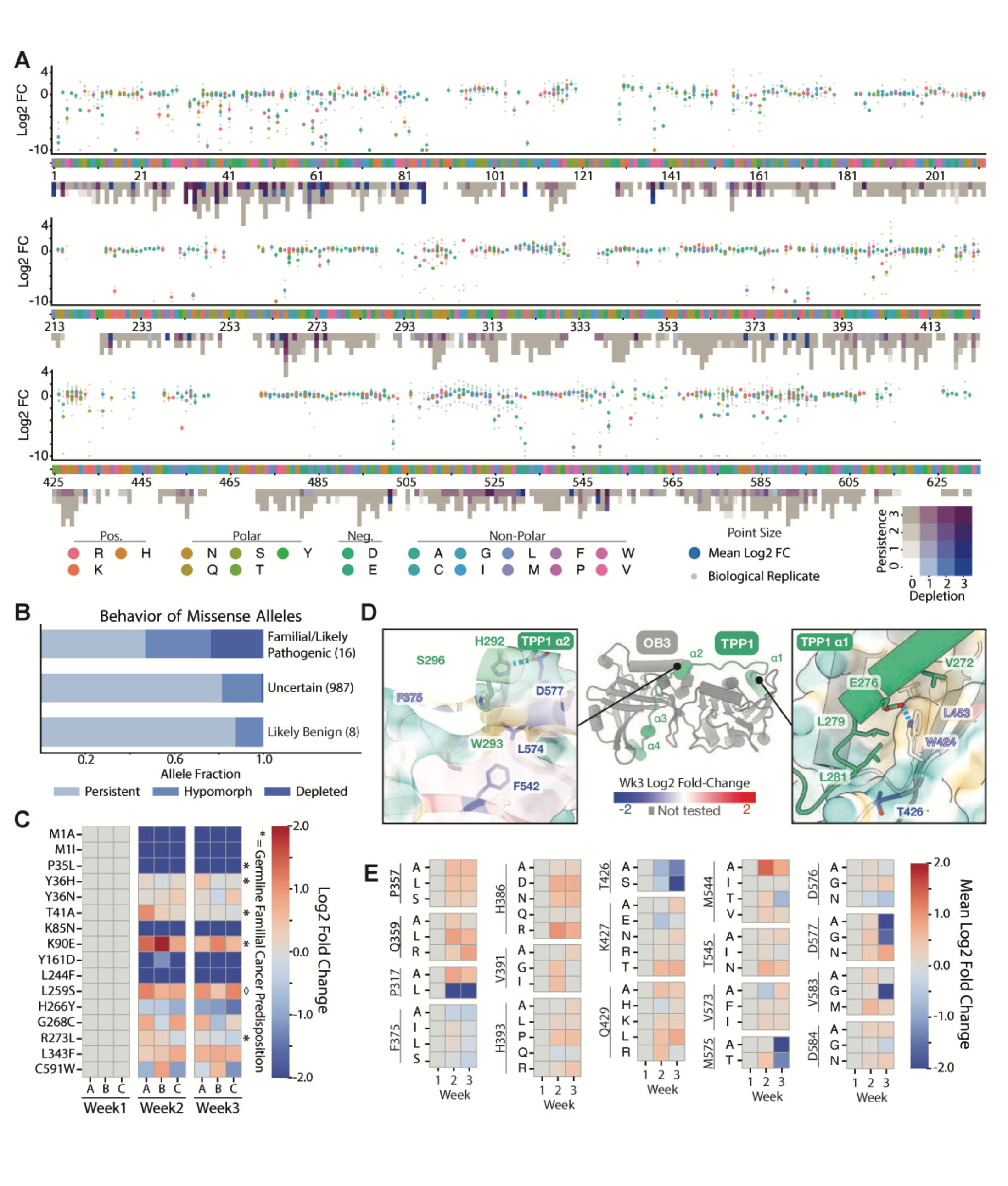
Many caPOT1 mutations are not simple loss-of-function mutations and instead fully support cellular proliferation. (**A**) Summary of the week 3 log2 fold-change of POT1 VUSs and all other substitution analyzed with the large dots indicating mean values at a given amino acid position and smaller dots showing individual biological replicates. Original amino acid identity is shown below the dot plot, with persistence/depletion scores for each substitution at a given position (see Materials and Methods section Data Analysis - Allele Classification). (**B**) Relative quantification of the proportion of alleles which are classified as persistent, hypomorphic, or depleted based on Wilcoxon rank-sum comparison to synonymous and frame-shift mutations occurring at similar week 1 allele frequencies (see Materials and Methods section Data Analysis - Allele Classification). Alleles are categorized as likely familial/likely pathogenic (based on ClinVar characterization, familial inheritance pattern, or disruption of the starting methionine), unknown, or likely benign (based on ClinVar characterization). (**C**) Heatmap showing log2 fold-change of alleles classified as “likely pathogenic” in panel B, with three biological replicates shown across three weeks of allele sampling. Asterisks indicate germline familial cancer predisposition mutations; diamond indicates a variant identified in familial idiopathic pulmonary fibrosis (*31*). (**D**) Structure of the TPP1 (green) POT1 (gray) interaction showing the four TPP1 alpha helices which contact POT1 (center). Insets show close views of TPP1 alpha helix 1 (ɑ1; right) and alpha helix 2 (ɑ2; left) contacting the hydrophobic surfaces of POT1. POT1 surfaces is colored by hydrophobicity, where yellow and teal represent hydrophobic or hydrophilic regions, respectively. POT1 residues are colored by log2 fold-change of alanine substitutions in that region. Model derived from PDB: 5UN7 (*44*). (**E**) Heatmaps showing mean log2 fold-change of three biological replicates across three weeks of sampling for the selected POT1 residues which contact TPP1 and were edited to multiple unique missense mutations. POT1 TPP1 contacts were identified by a Van der Walls overlap greater than or equal to −0.4 angstroms, ignoring interactions fewer than 4 bonds away (remaining loci with fewer unique missense mutations are shown in fig. S2E).

Surprisingly, a substantial subset of mutations that are either likely pathogenic (based on ClinVar classification) or are associated with an established familial and dominant inheritance pattern did not impact cellular proliferation (Fig. 3, B and C). One well-characterized familial mutation that falls into this class of mutations is R273L, which has been proposed to abrogate POT1’s electrostatic binding with ssDNA by disrupting interaction with the telomeric dT7 nucleotide (*5*, *22*, *46*, *49*). However, in our mutation scanning screen, both R273A and R273L were tolerated (Fig. 2E and Fig. 3C). Similarly, Q94 is another amino acid frequently mutated in melanoma and chronic lymphocytic leukemia and has been similarly shown to be essential for DNA binding (*5*, *24*, *33*, *45*). However, in our screen, both Q94H and Q94R persist (fig. S2D). Conversely, mutations which disrupt the starting methionine, or familial mutations proposed to dramatically disrupt protein conformation like P35L (*12*), do show strong depletion.

A similarly complex pattern arises when analyzing VUS which may impact POT1’s interactions with TPP1. Alanine substitutions at residues F542, L574 and D577, which contact residues of TPP1’s alpha helix 2 (*44*), deplete over the course of three weeks (Fig. 3D, left). However, interactions with TPP1’s alpha helix 1, including W424 and L453 do not show as strong of an effect (Fig. 3D, right). The cancer-associated C591W variant which was shown to indirectly disrupt POT1-TPP1 binding *in vitro*, likely through a conformational change (*33*, *44*), also did not have a significant effect on cell viability (Fig. 3C), nor did ClinVar VUS C591F, C591R, or C591Y (fig. S2E). A broader survey of amino acids shown to contact TPP1 supports the observation that only a small subset of mutations which could affect POT1 recruitment via TPP1 impact cellular viability (Fig. 3E and fig. S2E). In summary, these data demonstrate that only a fraction of caPOT1 mutations, including familial-inherited and molecularly-validated mutations, impaired cellular viability in our loHAP system over the course of three weeks. Together, these findings counter the hypothesis that all caPOT1 mutations are simple loss-of-function mutations and demonstrate that cellular viability of caPOT1 mutant cells in a hemizygous setting is not predictive for pathogenicity or cancer predisposition.

### POT1 VUSs can cause telomere elongation and trigger an ATR response without impairing cellular viability

If viability is not a reliable indicator of carcinogenic potential, we hypothesized that caPOT1 mutations may be better categorized by their effect on telomere length control, telomere protection or coordination of telomere C-strand binding. To test this, we first established the kinetics of DDR signaling at telomeres following acute POT1 mutagenesis by monitoring the accumulation of ɣ-H2AX telomere-dysfunction induced foci (TIFs) (*50*) over time following CRISPR/Cas9 mutagenesis with a single sgRNA (Fig. 4, A and B, and fig. S3, A and B). As expected, TIFs were detectable 72h after nucleofection in edited POT1 loHAP cells, but not in control cells. However, TIF burden surprisingly increased during the first week post editing and remained at a high level for three weeks, a time point where frame-shift mutations were largely absent from the mutant pool (Fig. 1, B and C, and fig. S1, B and C). In fact, a subset of TIF positive cells (conservatively defined as a minimum of 10 TIFs per cell) was detectable in screening mutagenesis pools generated for every exon of POT1 beyond three weeks in culture (Fig. 4C). To estimate the prevalence of alleles which induce DDR at telomeres without compromising cellular viability, we developed a high throughput genotyping and imaging analysis pipeline (Fig. 4D) to directly annotate TIF-causing mutations and estimate their prevalence by subpopulational sampling and Bayesian linear regression. Starting with arrayed subpools from mutagenesis of exons 8 and 9, this approach allowed us to identify several high-confidence TIF-causing mutations, including ClinVar VUSs L60R and S63I (Fig. 4E). Based on genotype complexity and TIF prevalence per well, our linear regression model estimates that more than 5% of all missense mutations present at week 3 in our mutagenesis pools are TIF positive.

**Fig. 4.**
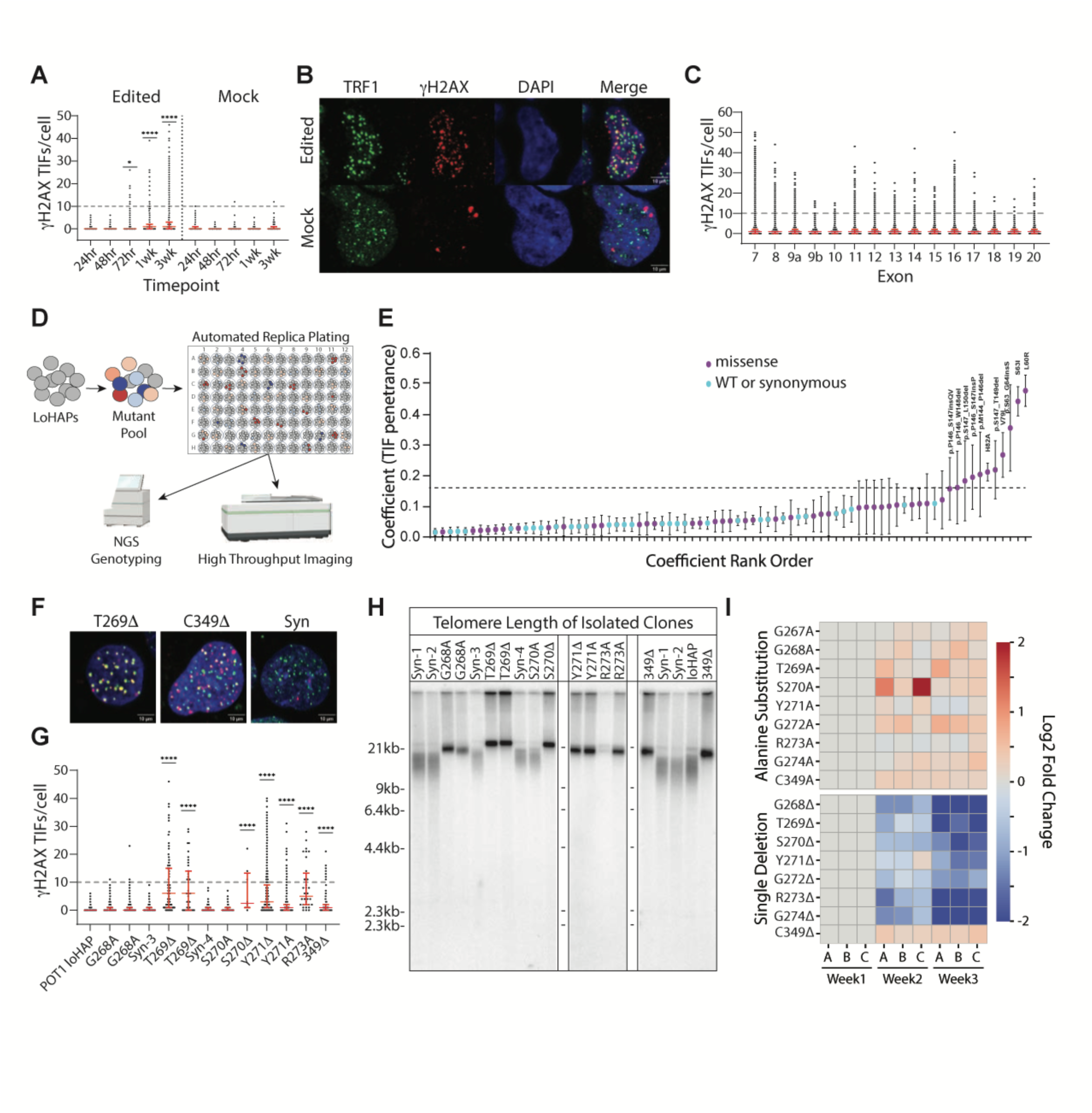
POT1 VUSs can cause telomere elongation without triggering an ATR response and without impairing cellular viability. (**A**) Quantification of γH2AX TIFs per cell over time in acute POT1 mutagenesis in loHAPs cells by a single guide RNA targeting exon 11, compared to unedited POT1 loHAPs. Red error bars indicate median + interquartile range. Fisher’s exact test with FDR correction for multiple comparison of nuclei with =>10 TIFs was done between Edited and Mock cells at each timepoint. * = p value <0.0332, ****= p value <0.0001. (**B**) Representative images for γH2AX TIFs characterized by the colocalization of TRF1 and γH2AX immunostaining signals in cells from acute POT1 mutagenesis compared to unedited POT1 loHAPs. Scale bar, 10 μm. (**C**) Quantification of γH2AX TIFs per cell at week 3 post mutagenesis in each exon showing mutants with >= 10 persistent γH2AX TIFs per cell were found across the entire POT1 gene. Red error bars indicate median + interquartile range. (**D**) Schematic of the high throughput imaging-based workflow to identify TIF-causing mutations. The POT1 mutant pools were arrayed out in 96-well plates by random subpopulation sampling followed by plate duplication, NGS genotyping and imaging. (**E**) The TIF+ coefficient of each POT1 mutant estimated in Bayesian regression model using datasets shown in panel D. Alleles were rank ordered by their coefficients. Error bars indicate SD. The coefficient threshold of calling TIF-causing alleles was defined by the highest confidence interval (mean+SD) of a synonymous variant. (**F**) Representative images for γH2AX TIFs in clonally isolated POT1 mutants. Scale bar, 10 μm. (**G**) Quantification of γH2AX TIFs per cell of clonally isolated POT1 mutants compared to unedited POT1 loHAPs. Duplicate genotypes indicate independently derived clones with the same mutation. Red error bars indicate median + interquartile range. Fisher’s exact test with FDR correction for multiple comparison of nuclei with =>10 TIFs was done between each cell line and POT1 loHAP. ****= p value <0.0001. (**H**) Telomere length analysis of clonally isolated POT1 mutant cell lines by Southern blot. Duplicate genotypes indicate independently derived clones with the same mutation. (**I**) Heatmap showing mean log2 fold-change of three biological replicates across three weeks of sampling for each indicated mutation. Alleles are grouped by mutation types.

The high prevalence of TIF-inducing alleles enabled us to clonally isolate TIF positive cells, including an allelic series of mutations proximal to the R273 residue mutated in familial cancer, mutations around S322, which is important for POT1s interaction with CTC1(*19*), as well as a clone with a single amino acid deletion of C349 proximal to POT1’s interaction domain with TPP1 (Fig. 4, F to I, and fig. S3, C to F). Mutant clones show a broad range of TIF phenotypes (Fig. 4, F and G, and fig. S3, C to E), which are not necessarily a direct predictor of telomere elongation (Fig. 4H and fig. S3F), or cell viability/proliferation in our initial screen (Fig. 4I). All assayed clones which show a robust, persistent DNA damage response at telomeres also have excessively elongated telomeres compared to synonymous mutants, including mutations affecting key residues such as R273 and S322. Consistent with previously published results (*19*, *20*), mutations affecting S322 also result in an overhang defect despite telomere elongation (fig. S3F). The strongest TIF responses, including T269Δ and Y271Δ also result in moderate allele depletion in our initial screen, even though these clones can be isolated and cultured beyond several months without a notable growth defect. However, even mutations such as G268A which do not show a significant TIF phenotype result in maximal telomere elongation without impacting cellular viability, establishing these mutations as true separation of function alleles in POT1.

### POT1 mutations combinatorially activate ATR-mediated ssDNA damage response signaling

The broad range of DDR phenotypes which persist in long-term culture suggest that instead of a binary telomere protection/deprotection phenotype, there may instead be a graded response to partial telomere deprotection which does not impair cell growth or viability until a cellular threshold is reached. To test this hypothesis and exclude the possibility that we have artificially selected for clones which are incapable of responding to ATR signaling, we performed a series of intragenic synthetic lethality experiments in which clones derived from the initial POT1 screen were re-edited at a secondary locus within the same POT1 allele using the same experimental approach as the initial screen (Fig. 5A). As we previously demonstrated, TIF-inducing mutations are well tolerated in loHAP cells or other TIF-negative cell lines such as the diploid parental hESCs or the S270A mutant clone; however, the same mutations show strong depletion in cells which already carry a TIF-inducing POT1 mutation such as C349Δ, R273A, or T269Δ (Fig. 5B and fig. S4). This intragenic synthetic lethality indicates that additional POT1 mutations can compound effects on cellular viability and that distinct domains of POT1 similarly activate the ATR-mediated single-stranded DNA damage response. This finding provides experimental evidence for previous predictions that the POT1-TPP1 heterodimer functions as a relay that can adopt continuous conformations which differentially expose the telomeric overhang (*46*).

**Fig. 5.**
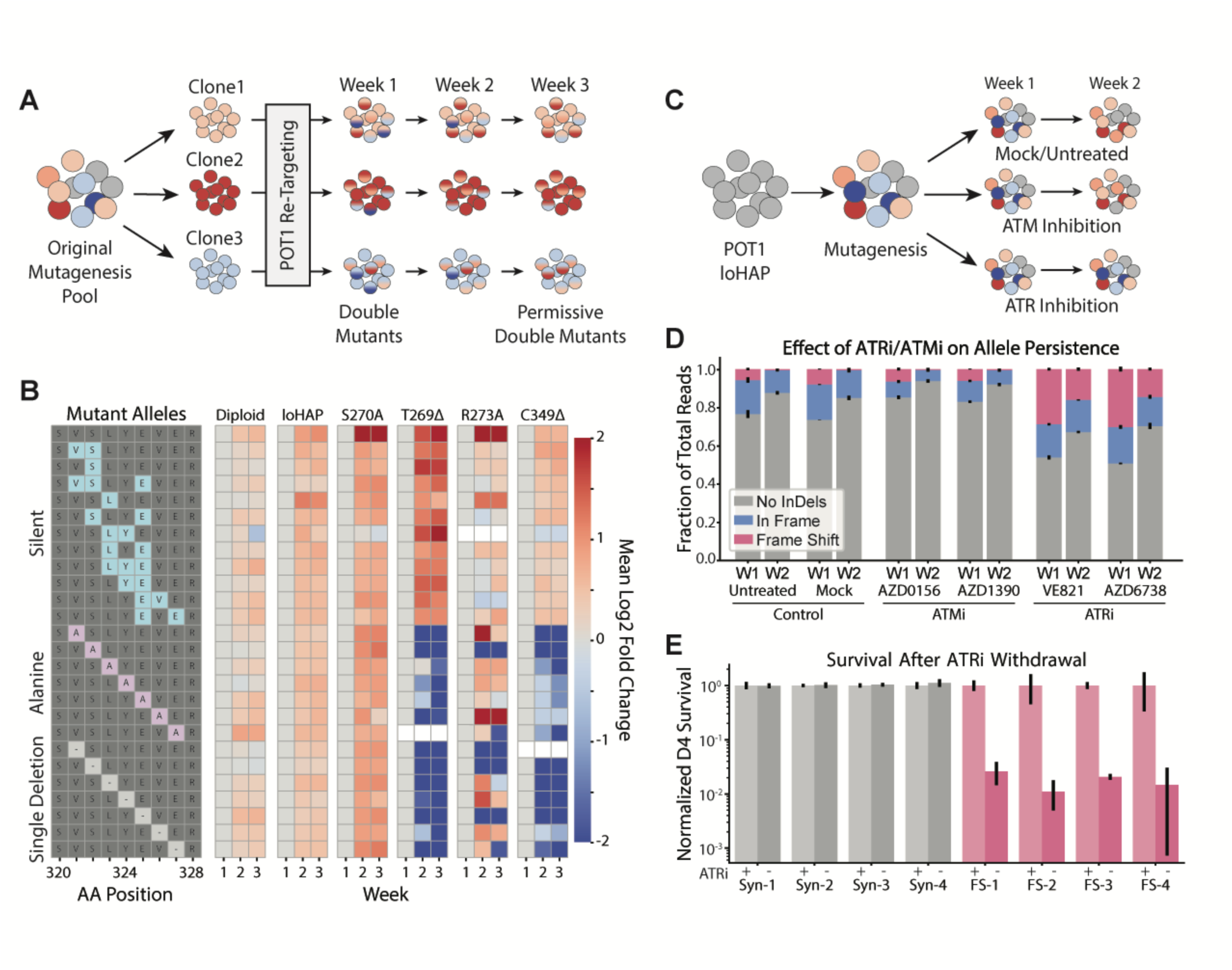
POT1 mutations drive cell lethality through ATR-mediated DNA damage response pathway activation. (**A**) Schematic of intragenic synthetic lethality experiment: Clones isolated from the initial POT1 screen were re-targeted to generate secondary mutations within the same POT1 allele which were then assayed over three weeks. (**B**) Heatmap showing mean log2 fold-change of allele frequency of three biological replicates across three weeks of sampling for each indicated cell lines (diploid wildtype, loHAPs, S270A, T269D, R273A, and C349D). Alleles introduced by the POT1 retargeting are grouped by category: silent, alanine substitution, or single amino acid deletion. Alleles which were not detected in all three biological replicates in an indicated cell line are shown in white. (**C**) Schematic of ATR/ATM inhibition screen. ATMi/ATRi was added immediately following sgRNA delivery. (**D**) Relative quantification of CRISPR/Cas9-mediated insertion and deletion alleles for two weeks post sgRNA delivery and drug treatment; three biological replicates, with error bars representing the standard error of the mean. (**E**) Quantification of cell survival four days post ATRi withdrawal (-) compared to continued maintenance on ATRi (+) for four frame-shift clones (pink) and four synonymous mutant clones (gray). Error bars show the standard error of the mean.

If cell viability in POT1 mutant clones is determined by a threshold of DDR signaling, we hypothesized that dampening this signaling pathway may allow more deleterious alleles to persist in culture (Fig. 5C). Indeed, only ATR inhibition (ATRi) and not ATM inhibition (ATMi) increased the persistence of frame-shift alleles past two weeks in bulk culture, confirmed by two independent inhibitors for each pathway (Fig. 5D). Surprisingly, despite the continued depletion of these alleles in bulk, continuous ATRi enabled clonal isolation of frame-shift mutants from all tested POT1 protein domains, encompassing five different exons and spanning all three POT1 OB folds (table S2). These clones are stable in culture for over 6 months provided ATRi is maintained. However, withdrawal of ATRi results in ∼100-fold reduction in cell viability in four days (Fig. 5E), indicating that the persistence and viability of these POT1 loss-of-function clones under ATRi is not a result of clonal adaptation or ATR pathway silencing. Therefore, the only role of POT1 which is essential for cell survival is the suppression of ATR signaling.

### caPOT1 mutants show qualitatively and quantitatively different TIF phenotypes than full loss of POT1

To test whether all consequences of POT1 mutations could be fully repressed by ATRi, we assayed DDR phenotypes and telomere elongation in frame-shift clones alongside T269Δ and C349Δ. Even under ATRi, POT1 frame-shift clones still exhibit γH2AX TIFs, indicating that our inhibitors may be insufficient to completely dampen a highly activated ATR signaling pathway. However, 48-hour ATRi withdrawal significantly increased the TIF burden in POT1 frame-shift clones (Fig. 6A). As an orthogonal approach, we also performed a targeted intergenic knockout screen in T269Δ and C349Δ mutant cell lines by knocking out ATR and 12 other DDR genes. We then stained for TIFs and assayed which gene knockouts reduced TIF burden in targeted cells. In both cell lines, ATR, RPA2, TOPBP1 and RAD51 knockouts were consistently among the genes leading to the greatest suppression in TIFs (fig. S5, G and H), suggesting that POT1-induced TIF phenotypes are truly ATR dependent.

**Fig. 6.**
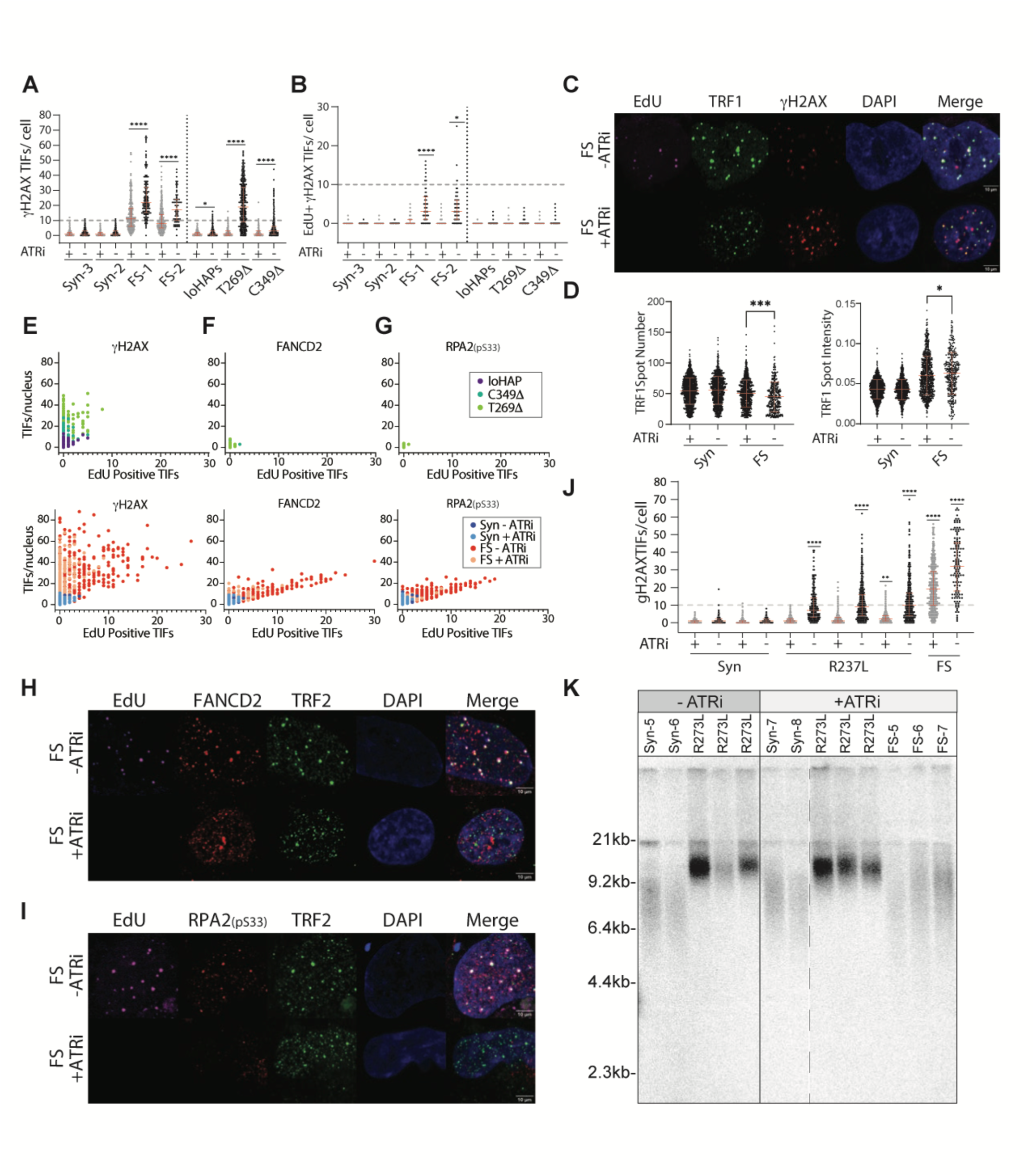
CaPOT1 mutations elongate telomeres independently from ATR signaling and have quantitatively and qualitatively different ATR-mediated TIFs compared to frame-shift clones. (**A**) Quantification of γH2AX TIFs per cell under ATR inhibition and 48 hours post ATRi withdrawal. Red error bars indicate median and interquartile range. Fisher’s exact test with FDR correction for multiple comparison of nuclei with =>10 TIFs was done between ATRi + and - conditions for each cell line. * = p value <0.0332, ****= p value <0.0001. **(B)** Quantification of γH2AX TIFs that colocalize with sites of EdU incorporation outside of S-phase under ATR inhibition and 48 hours post ATRi withdrawal. Red error bars indicate median + interquartile range. Fisher’s exact test with FDR correction for multiple comparison of nuclei with =>10 EdU positive TIFs was done between frame-shift (FS) and synonymous (Syn) cells and between mutant and POT1 loHAP cells. * = p value <0.0332, ****= p value <0.0001. (**C**) Representative images of EdU incorporation outside of S-phase and γH2AX TIF colocalization outside of S-phase in POT1 frame-shift cells ± ATRi. Scale bar, 10μm **(D)** TRF1 Spot Number and TRF1 Spot Intensity plots. Data is representative of four synonymous and four frame-shift clones ± ATRi. Red error bars indicate means + SD. One-way ANOVA using Bonferroni’s multiple comparison test was used to indicate significance. * p value <=0.0332, *** p value <=0.0002. **(E)** Frequency plots for sites of EdU positive γH2AX TIFs (x-axis) vs. total γH2AX TIFs per cell (y-axis). The top panel shows the frequency of distribution in loHAP, C349D and T269D POT1 mutant cells; the bottom panel shows frame-shift and synonymous mutations with and without ATRi. **(F)** Frequency plots as in E for the colocalization of EdU positive FANCD2 TIFs vs. total FANCD2 TIFs per cell. **(G)** Frequency plots as in E for the colocalization of EdU positive RPA2 (pS33) TIFs vs. total RPA2 (pS33) TIFs per cell. **(H)** Representative images for EdU and FANCD2 TIF colocalization in POT1 frame-shift cells ± ATRi. Scale bar, 10 μm **(I)** Representative images for EdU and RPA2 (pS33) TIF colocalization in POT1 frame-shift cells ± ATRi. Scale bar, 10 μm **(J)** Quantification of γH2AX TIFs for mutants derived under ATRi. Withdrawal of ATRi is indicated below the x-axis. Red error bars indicate median + interquartile range. Fisher’s exact test with FDR correction for multiple comparison of nuclei with =>10 TIFs was done relative to Syn - ATRi cells. ** = p value <0.0021, ****= p value <0.0001. **(K)** Telomere length analysis of R273L, synonymous (Syn) and frame-shift (FS) clones derived either under ATRi or without ATRi. Duplicate R273L genotypes indicate independently derived clones, whereas Syn and FS clones are labeled according to table S2.

Since previous experiments have shown activation of HDR machinery at telomere ends following POT1 depletion (*16*), we assayed EdU incorporation at telomeres outside of S-phase as a proxy for HDR mediated DNA synthesis. Colocalization of EdU foci with γH2AX TIFs is seen exclusively in POT1 frame-shift cells after ATRi withdrawal (Fig. 6, B and C), despite the strong γH2AX TIF phenotype in frame-shift clones under ATRi. The appearance of EdU+ TIFs in frame-shift clones post ATRi withdrawal correlates with a reduction in the number of telomere foci detected by TRF1 staining, but an overall higher spot intensity (Fig. 6, C and D), supporting the hypothesis that EdU incorporation is likely due to telomere clustering and recombination.

This suggests that loss-of-function mutants under ATRi recognize unprotected telomeres with early γH2AX accumulation, but that full ATR pathway activation is required for downstream telomere effects. To test this, we assayed FANCD2, RPA2 (pS33) and 53BP1 colocalization with EdU+ TIFs in each cell line (Fig. 6, E to I, and fig. S5, A and B). Indeed, frame-shift clones show 53BP1 positive TIFs under ATRi, but FANCD2 and RPA2 (pS33) only accumulate following ATRi withdrawal and strongly correlate with EdU incorporation, demonstrating that TIFs following ATRi withdrawal are both qualitatively and quantitatively different from those observed under ATRi. RPA2 is known to bind exposed ssDNA at the telomere in the absence of POT1, where ATR signaling leads to phosphorylation of RPA2 Serine 33 (*34*, *51*). However, even in frame-shift mutants, unphosphorylated RPA2 was not highly detected at telomeres until ATRi withdrawal (fig. S5, C and D).

Conversely, neither ATRi-withdrawal phenotype – EdU incorporation outside of S phase nor accumulation of FANCD2 and RPA2 (pS33) – can be detected in the T269Δ and C349Δ clones regardless of ATRi condition (Fig. 6, B and E to G). Further, ATRi abrogates the γH2AX TIF phenotype to background levels indistinguishable from the parental loHAP clones (Fig. 6A). Therefore, these clones represent a sub-activated state of ATR signaling wherein telomeres accumulate γH2AX and 53BP1 but fail to fully activate the ATR signaling cascade which would result in RPA2 phosphorylation and ultimately drive cell lethality.

### ATR signaling is not required for telomere elongation in caPOT1 mutants

Despite the ability of ATRi to repress all assayed DNA damage response phenotypes, ATRi does not prevent telomere elongation. Derivation of R273L or C349Δ clones with and without ATRi both result in robust telomere elongation (Fig. 6, J and K, and fig. S5, E and F). However, despite showing a strong TIF phenotype, frame-shift mutants at the same locus exhibit shorter telomeres than the R273L clones at time of assay (∼6 weeks post-editing).

If both telomere elongation and DDR signaling were simply a function of telomere deprotection, or if caPOT1 mutations merely compromised protein stability, frame-shift mutations should show the strongest degree of both phenotypes. Therefore, the rapid telomere elongation in the caPOT1 mutants compared to the frame-shift mutants can only be explained by three possibilities: either residual POT1 function is required for rapid telomere elongation, caPOT1 mutations gain *de novo* stimulatory effects on telomerase, or elongation is actively counteracted in frame-shift clones, perhaps through nucleolytic degradation of unprotected telomeres. Together, these data demonstrate that subset of caPOT1 mutations are separation of function mutations which do not fully deprotect telomeres. Instead, they elicit a sub-threshold DDR phenotype which is insufficient to fully activate ATR and these mutations drive telomere elongation in an ATR-independent manner.

### Loss of POT1 does not impair telomere replication

The simplest explanation for the decreased rate of telomere elongation in the frame-shift clones is that loss-of-function mutations result in an inability to properly replicate telomeres, since POT1 is thought to limit excessive 5’ resection and recruit the CST complex to enable fill-in synthesis of the lagging strand (*20*, *45*, *52*). CTC1 loss was previously shown to cause decreased cell viability as a function of gradual telomere loss (*53*). To confirm this phenotype in our hESC, we engineered conditional CTC1^F/+^ and CTC1^F/-^ hESC lines such that induction of Cre recombination results in either CTC1^-/+^ heterozygous or CTC1^-/-^ null cells (fig. S6A). As expected, the loss of CTC1 led to a rapid increase in TIFs and an eventual loss of cell viability at ∼24 days post loop-out of the conditional allele, concurrent with telomere shortening and overhang extension (Fig. 7A and fig. S6, B to D).

**Fig. 7.**
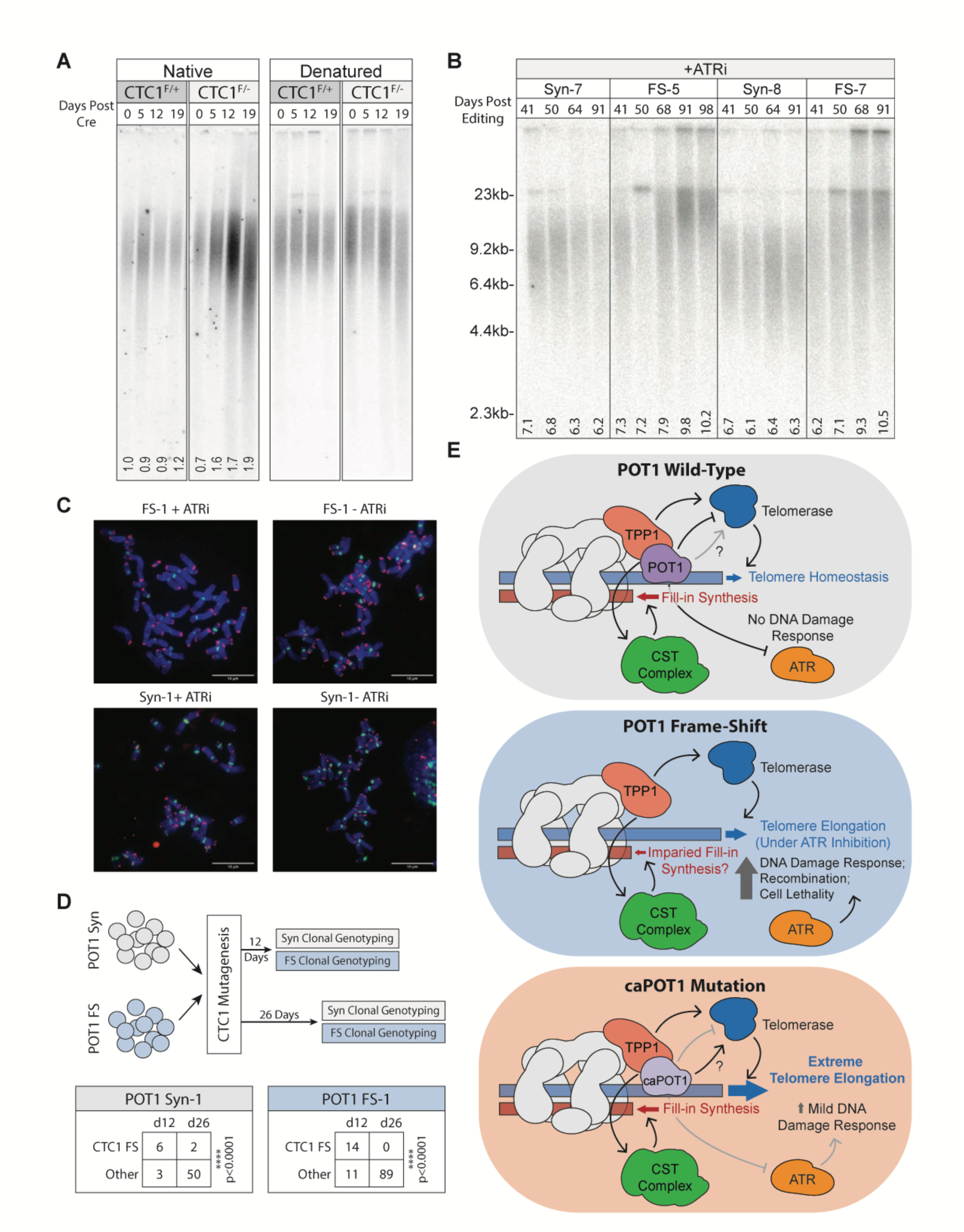
Under ATRi, POT1 is not essential for telomere replication or maintenance in hESC. **(A)** Telomere length analysis of conditional CTC1^F/+^ and CTC1^F/-^ hESC cells before and after induction of Cre with 2µM 4-hydroxytamoxifen under native (left) and denaturing (right) conditions. Relative overhang intensity normalized to lane 1 (CTC1^F/+^ day 0) is indicated under the native gel. Telomere probe: (CCCTAA)_3_ **(B)** Time-course telomere length analysis of two synonymous (Syn-7 and Syn-8) and two frame-shift (FS-5 and FS-7) clones spanning days 41-98 post nucleofection. **(C)** Representative metaphase spreads for frame-shift (FS-1) and synonymous (Syn-1) cells under ATRi inhibition as well as after ATRi withdrawal. Centromere stain is shown in green whereas telomeres are shown in red. Scale bar, 10 μm. Results are quantified in table S3. **(D)** CTC1 targeting experiment in POT1 Syn-1 and POT1 FS-1 cells. POT1 Syn-1 and FS-1 cells were nucleofected as described in the Materials and Methods section, and individual clones were genotyped at 12 days and 26 days post nucleofection. “CTC1 FS” refers to clones which carry homozygous or compound heterozygous frame-shift mutations in CTC1. “Other” refers to any clone which carries at least one unedited or in-frame edited allele. Significance was determined by Fisher Exact test. **(E)** Model of POT1 interactions and telomere end homeostasis in a POT1 haploid setting. Under wild-type conditions (top), POT1 represses ATR-mediated DDR and telomere length homeostasis is maintained. In POT1 frame-shift mutations (middle), ATR signaling is highly activated and results in cell death unless inhibited by ATRi; fill-in synthesis occurs but may be dysregulated and telomeres elongate. In the case of caPOT1 mutations (bottom), ATR signaling may be partially activated, but DNA damage responses are mild; POT1 repression of telomerase activity is diminished, and fill-in synthesis is unaffected, so telomeres elongate dramatically and rapidly.

In contrast, frame-shift clones under constant ATRi continuously elongate their telomeres over the course of several months (Fig. 7B). However, even after 13 weeks in culture they do not recapitulate the uniform elongation and narrow window of telomere lengths of caPOT1 mutations at the same locus (Fig. 6K), but instead show much more heterogeneous telomeres. This long-term telomere elongation in frame-shift mutants is consistent across 6 independently derived clones compared to time-matched synonymous mutants (Fig. 7B and fig. S7A). Even after ATRi withdrawal, loss-of-function cells retain long telomeres up to 72 hours with no catastrophic telomere shortening or overt telomeric defects prior to cell death (Fig. 7C and fig. S7A). Despite a slight increase in sister-chromatid fusions and fragile telomeres, frame-shift cells under ATRi also appeared remarkably devoid of excessive telomeric abnormalities as detected by metaphase spreads and do not show overt overhang abnormalities until ATRi is withdrawn (Fig. 7C, fig. S7, C and D, and table S3). Although frame-shift clones show a dramatic decrease in total metaphases, they otherwise show fewer abnormalities than viable frame-shift cells under ATRi.

These data support two conclusions: First, since frame-shift clones under ATRi do not show overt replication stress, the genomic instability triggered by loss of POT1 is caused by ATR-mediated DDR signaling, not by inherent replication stress due to unprotected telomeres. Second, POT1 is not essential for fill-in synthesis. Recent *in vitro* studies have shown that priming for lagging-strand DNA synthesis cannot occur at the telomeric overhang, and requires CST to counteract lagging-strand shortening (*52*). Under this assumption, extension of the 3’ overhang via telomerase would be insufficient to rescue shortening of the double-stranded portion of the telomere and prevent cell death, indicating that fill-in synthesis must still be accomplished in our frame-shift clones under ATRi despite loss of POT1. Indeed, CTC1 is still essential in POT1 frame-shift clones, as CTC1 knockout in both POT1 frame-shift or synonymous clones are viable at day 14 but show dramatically reduced colony survival after 26 days in culture, as measured by both total clones detected and cell counts per clone (Fig. 7E and fig. S7B).

Therefore, the only essential function of POT1 in our hESC system is to repress ATR signaling and POT1 is not required for telomere replication or fill-in synthesis in the absence of ATR signaling. This suggests a model wherein wild-type POT1 regulates telomere length homeostasis by controlling access of telomerase to the telomere end and repressing ATR-mediated DDR signaling (Fig. 7E, top). On the other hand, loss of POT1 caused by frame-shift mutations results in catastrophic DNA damage signaling, culminating in cellular lethality. If ATR signaling is repressed, telomerase has greater access to the 3’ telomeric overhang, resulting in telomere elongation. For this elongation, fill-in synthesis must still occur, though it remains possible that this process is somewhat impaired without POT1-mediated regulation since telomere elongation is not uniform and telomeres become heterogeneous (Fig. 7E, middle). However, some caPOT1 mutations reduce the ability of POT1 to repress telomerase activity and may in fact stimulate telomere elongation through an unknown mechanism while retaining the ability to regulate fill-in synthesis and maintain ATR-mediated DNA damage response signaling beneath a cellular threshold which would trigger telomeric recombination or cell death. Together, this results in rapid and dramatic telomere elongation (Fig. 7E, bottom).

## Discussion

In this study, we assayed over 600 clinically relevant variants of unknown significance and used single-codon deletion and alanine scanning to functionally characterize POT1 alleles relevant to human disease. As our POT1 loHAP system provides a primary human cell system with a stable genome and intact telomere maintenance and DDR pathways, these mutations closely parallel the early stages of carcinogenesis. The presence of inherited early truncating mutations and substitution of the starting methionine in POT1 cancer spectra indicate that full loss-of-function alleles drive cancer through POT1 haploinsufficiency. These alleles strongly depleted in our screen with a median fold-change of 2^-9.97^ demonstrating the high dynamic range and sensitivity of our cell system. However, other caPOT1 mutations – including validated familial mutations – show less depletion and a broad array of phenotypes which culminate in varied DNA damage responses and telomere elongation without affecting cellular viability. Thus, allele depletion in our model is not a deterministic predictor of human pathogenicity and cancer-associated phenotypes do not scale linearly with cell viability but are functionally independent. Instead, mutations throughout POT1 converge on a common phenotype of robust telomere elongation, supporting the hypothesis that the flexible interaction between POT1-TPP1 and telomeric ssDNA may integrate distal conformational changes, resulting in varying degrees of telomeric overhang exposure (*46*, *54*).

This also explains the broad range of tolerated γH2AX TIF phenotypes across caPOT1 mutations and reveals a “Goldilocks zone” where DDR signaling is insufficient to affect cell viability while increased telomerase accessibility still has a large relative effect on telomere length. This suggests that the primary constraint on POT1 sequence conservation may be telomere length homeostasis rather than telomere end protection.

The initial scope of our work was performed under the hypothesis that all caPOT1 alleles which drive cancer are loss-of-function alleles. Our results clearly demonstrate that even under hemizygous conditions, caPOT1 alleles can be separated and classified based on cellular phenotypes of viability, telomere elongation, and DDR signaling. We have previously demonstrated that a set of heterozygous caPOT1 mutations are capable of telomere elongation without driving a detectable DNA damage response (*27*). Because telomere elongation is the more penetrant phenotype in a heterozygous setting and, elongation occurs independently of ATR signaling, we hypothesize that the remaining wild-type allele may dampen DDR signaling while preserving telomere elongation.

Lastly, we identified a class of separation-of-function caPOT1 alleles which outperform frame-shift alleles in rapid telomere elongation while maintaining DDR signaling below a cell-intrinsic threshold for full pathway activation of ATR. This class of caPOT1 mutations retains or gains telomere length stimulatory roles which are not present in full loss-of-function alleles, identifying them as a class of oncogenic driver mutations rather than loss of function alleles. We propose three potential mechanisms for this elongating activity:

First, residual POT1 interaction with CST may aid rapid telomere elongation. While we demonstrate that POT1 is not essential for telomere fill-in synthesis during DNA replication in our cells, it remains possible that POT1-CST interactions modulate the frequency or activity of C-strand synthesis. Therefore, inefficient CST recruitment may result in a mild degree of overhang extension which is sporadically corrected by association of CST with the telomere through other means, perhaps via interaction with TPP1 (*28*).

Second, deprotection of telomeres in POT1 frame-shift mutants under ATRi may result in telomere degradation. Unless another ssDNA binding protein protects the overhang, the free ssDNA end may be accessible to exonuclease processing, resulting in slower telomere elongation since the activity of telomerase is required to counteract this active degradation. Alternatively, deprotection of the single-stranded 3’ overhang may compromise t-loop formation and result in catastrophic telomere loss, perhaps resulting in the heterogenous telomeres we detect in our frame-shift mutants.

However, the lack of an overt telomeric overhang phenotype in POT1 frame-shift clones under ATRi leads us to favor a third model: caPOT1 mutations may have direct stimulatory effects on telomerase. The POT1-TPP1 heterodimer has been shown to increase telomerase repeat addition *in vitro* (*55–57*). While the TPP1-TERT interaction is well characterized, the recently solved structure of telomerase bound to shelterin also suggests that POT1 directly contacts the TERT TEN domain (*49*, *58*), and disruptions to this interaction domain reduce telomerase elongation (*59*). Thus POT1 may also function as a positive regulator of telomere length, though the precise mechanisms remain unclear. Additional experimentation would be required to determine whether caPOT1 mutations merely retain this stimulatory activity despite increased accessibility to the 3’ overhang, or whether these represent *de novo* stimulatory effects, perhaps explaining why heterozygous mutations act dominantly in cancer development.

## Acknowledgments

We dedicate this work to our friend and colleague John Boyle, who passed away on August 9th, 2024, in honor of his personal and professional battle against cancer and scientific contribution to this study. The AAVS1 targeting construct to express CRE-ER was a generous gift from Frank Soldner, Albert Einstein Institute of Medicine.

## Funding

American Cancer Society grant 133396-RSG-19-029-01-DMC (DH) Siebel Stem Cell Institute grant NIH R01-HL131744 (DH)

CIRM training program EDUC4-12790 (JS)

Jane Coffin Childs Postdoctoral Fellowship (GEG)

NIH grant R35-GM134922 (YSS)

## Author contributions

AM, JS, RB, HL and DH designed the experiments with the advice of YSS; experiments were performed by JS, RB, HL and DH with the help of ST, RL, RH, and AP; the TIF linear regression model was developed by JG. and HL with the advice of YSS and MJ; SJ and JB generated the conditional CTC1 KO hESCs; data was analyzed by AM, JS, HL and DH; structural analysis and visualization was performed by GEG.; AM, JS, HL and DH.wrote the paper.

## Materials and Methods

### Human Embryonic Stem Cell Culture

Pluripotent stem cell research is approved under 2012-12-024 by the Stem Cell Research Oversight Committee at the University of California, Berkeley. WIBR3 hESCs,(*61*) National Institutes of Health stem cell registry #0079) were cultured on 4.1 × 10^5^ cm^−2^ irradiated mouse embryonic fibroblasts (MEFs) in hPSC medium (Dulbecco’s modified Eagle medium/Nutrient Mixture F-12 (DMEM/F12), 20% KnockOut Serum Replacement, 1× Non-Essential Amino Acids (NEAA), 1 mM glutamine, 1× penicillin/streptomycin, 0.1 mM β-mercaptoethanol and 4 ng/ml heat-stable basic fibroblast growth factor). Media was changed daily and cells were passaged weekly with 1 mg/mL collagenase IV.

During screening, cells were cultured feeder-free using matrigel-coated culture dishes and MEF-conditioned hSPC media (Dulbecco’s modified Eagle medium/Nutrient Mixture F-12 (DMEM/F12), 20% KnockOut Serum Replacement, 1× Non-Essential Amino Acids (NEAA), 1 mM glutamine, 1× penicillin/streptomycin, and 0.1 mM β-mercaptoethanol was plated onto 4.1 × 10^5^ cm^−2^ irradiated MEFs for 24 hours, harvested, supplemented with 10 ng/ml heat-stable basic fibroblast growth factor). Media was changed every other day. Cells were single-cell passaged weekly using Trypsin 0.25% EDTA, which was inactivated using 5% Fetal Bovine Serum (FBS) added to the hPSC medium. The day before and after passage, medium was supplemented with 10 µM Y27632 to increase cell survival.

### Nucleofection of hESCs

hESCs cultured on MEFs were detached from feeder cells by treating with 1 mg/ml collagenase IV for 20-30 min. Colonies were then suspended using hSPC Wash Media (DMEM/F12, 5% FBS, 1x penicillin/streptomycin), then sedimented twice for 5 min at room temperature (RT) to remove MEF carry-over. Following aspiration of the Wash Media and wash with phosphate-buffered saline, cells were dissociated to single cells by incubation with accutase for 5 min at 37C. Dissociated cells were washed with 10x volume of Wash Media, pelleted, resuspended in phosphate-buffered saline (PBS), and counted.

Nucleofection was performed as described in Li et al. 2023.(*43*) Briefly, 0.5-1 million hESCs were pelleted and resuspended in 20ul Lonza P3 nucleofection reagent, mixed with pre-assembled Cas9/sgRNA RNP with or without 100 pmol ssDNA oligo pool HDR donor templates, and nucleofected using program CA137 on Lonza nucleofector 4D.

Gene KO experiments were performed using multiguide Gene knockout Kit v2 (Synthego) targeting the specific genes of interest. Nucleofections were performed as described above. Cells were assayed for TIFs 7 days post nucleofection.

### LoHAP Generation

POT1 loHAPs were generated as described in Li et al. 2023.(*43*) Briefly, sgRNAs were chosen based on specificity scores(*62*, *63*) and located between 50-300bp downstream of the transcriptional end sites of POT1 and GRM8. After nucleofection, cells are seeded onto 96-well plates at 1,000 cells/well, or 100 cells per well. After 12 days of outgrowth, cells were replica plated and remaining cells were added to an equal volume of 2x PCR-compatible lysis buffer (100mM KCl, 4mM MgCl2, 0.9% NP-40, 0.9% Tween-20, 500µg/ml proteinase K, in 20mM Tris pH 8). The cell lysate was incubated overnight at 50C, then heated for 10 min at 95C to inactivate proteinase K. Cell clones were genotyped using sanger sequencing of a junction-spanning PCR (PCR primers outside of the intended deletion) and NGS of a SNP located intronically between exon 15 and 16 (HG38 chr7:124837128 A/G). Primer sequences included in Supplemental Table 4. PCR was performed using either Titan or PrimeStarGXL polymerases using 2uL of the lysate. Wells of interest with a single nucleotide remaining at a SNP were then further subcloned by low-density seeding or manual picking to ensure clonality.

### Deep Scanning Mutagenesis

Mutagenesis was performed as described in Li et al 2023,(*43*) with minor adjustments. Targeted designer mutations were introduced with pools of 150–200 nt single-stranded oligonucleotide HDR templates centered at the CRISPR/Cas9 cut site. Oligo pools were purchased from Integrated DNA Technologies (IDT). HDR templates were designed with synonymous CRISPR/Cas9 blocking mutations to increase integration efficiency; when synonymous PAM-ablating mutations were not feasible, sgRNA-blocking mutations were introduced proximal to the cut site. All HDR templates contain a ≥2 nucleotide change relative to the wild-type sequence to distinguish designer mutations from sequencing error.

Mutagenesis was performed by nucleofection into 1 million loHAP cells as described above with the inclusion of 100pmol of ssDNA oligo pool HDR donor templates. Immediately following nucleofection, cells were pooled based on sgRNA cutsite location into pools which could be interrogated by a single sequencing amplicon (141 total sgRNAs divided into 16 pools), then divided into three biological replicates derived from the same nucleofection event. Each biological replicate was cultured as described above, and single-cell passaged weekly at a ratio of 1:20. Remaining cells were split between two frozen cell stocks and two lysis tubes as described under loHAP generation.

Sequencing was performed by PCR amplification using TAKARA GXL polymerase and the cell lysis equivalent of between 10,000-20,000 genomes was added per technical replicate. Between 8 and 24 technical replicates were performed per biological replicate until the number of unique alleles detected was saturated. Amplicons were purified using SPRI bead purification at the UC Berkeley DNA Sequencing Facility, i5/i7 barcoded, pooled, and run on 150 PE iSEQ for quality control before NextSeq or NovaSeq X deeper sequencing at the Center for Advanced Technology in UCSF. All sequencing primers, HDR oligo templates, and sgRNA sequences are contained in tables S4, S5, and S6, respectively.

### Frame-shift Analysis

For single-sgRNA experiments, NGS fastq files were analyzed with CRISPResso2(*64*) to merge reads, quality filter, and separate alleles. Frameshift_analysis.txt outputs from CRISPResso2 were visualized into barplots using python library matplotlib.

### Data Analysis - Allele Classification

For pooled sgRNA mutagenesis experiments, demultiplexed NGS fastq files from each technical replicate were initially analyzed using CRISPResso2(*64*) to merge reads, quality filter, and separate alleles per technical replicates. Allele frequency tables were then read into a custom python script for further allele classification. First, alleles which showed full HDR (defined by complete integration of the HDR template) or partial HDR (defined by integration of the HDR template to one side of the predicted cutsite) were identified. Then, cutsite positions for all sgRNAs in the amplicon were used to define editing windows +/- 2nt from any cutsite position. Lesions not extending from these cutsite positions were filtered out and any single nucleotide substitutions not at cutsites and not generated by HDR were determined to be PCR error and corrected to wild-type. Any alleles showing only a single nucleotide change (insertion, deletion or substitution), alleles which are PCR corrected to wild-type, alleles showing multiple editing events, and alleles containing insertions were excluded from the analysis. Remaining alleles were translated *in-silico* and collapsed based on proposed protein changes.

Following allele classification, technical replicates were summed per amplicon and only alleles with an overall allele frequency >1×10^-6^ were retained. Alleles were then normalized to read depth to generate allele frequencies and then their respective week 1 samples to calculate fold-change. To calculate log2 fold-change and enable visualization of alleles which completely deplete (and thus have a fold-change of 0), 0.001 was added to all fold-change values before applying the log2. Heatmaps, dot plots, and other visualizations were generated using matplotlib and seaborn.

### Alleles were then scored based on two metrics

#### Persistence/Depletion Scoring

To account for increased variance at lower allele frequencies, Log2 fold-change for each biological replicate was plotted against week 1 allele frequency and the 95th or 5th percentile was calculated for frame-shifts or synonymous mutants respectively (Shown in Supplemental Figure 1F), using a scanning window of 180 data points. Curves were fit to these percentile graphs using the scipy optimize.curve_fit module. Alleles were then classified based on these curves: For each biological replicate falling below the curve defined by the synonymous mutations, a Depletion Score of +1 was added, to a maximum of three (three biological replicates). Similarly, replicates above the frame-shift curve were awarded a Persistence Score of +1, to a maximum of 3.

#### Statistical Comparison

All alleles present in at least 2 biological replicates were statistically compared to the 180 synonymous and frame-shift alleles closest to the same week 1 allele frequency. As frame-shift alleles do not show a normal distribution, Wilcoxon rank sum was used to determine if the allele falls significantly above the frame-shift distribution and significantly below the synonymous distribution. Alleles which were not significantly depleted compared to the synonymous mutations were classified as Persistent. Alleles which were not significantly enriched compared to the frame-shift alleles were classified as Depleted. Alleles significantly different from both were classified as Hypomorphs.

### Structural Visualization

A python script was used to map log2 fold-change in allele frequency onto the PDB file B-factor column. Structural predictions were performed using AlphaFold2 through a local installation of Colabfold 1.2.0,(*65*) running MMseqs2(*66*) for homology searches and AlphaFold2-Multimer(*67*) for the prediction. Models were visualized in ChimeraX.(*68*) PDB IDs are included in the figure legends.

### TIF Staining

Cells were plated on Matrigel (Corning) coated Phenoplate-96 imaging plates (Revvity) at a seeding density of ∼20 000 cells per well 24hr prior to fixation with 4% Paraformaldehyde for 10 min at RT. Following a wash in 1x PBS, cells were permeabilized in 0.1% Triton X-100 for 15 min at RT, before an additional wash in 1x PBS and 15 min blocking in Blocking buffer (3% BSA + 5% horse serum and 0.1% Tween-20 in PBS) at RT. Primary (anti-TRF1: 1/10 000, anti-TRF2: 1/2000, anti-gH2AX: 1/2000, anti-53BP1: 1/2000, anti-RPA32 s33: 1/2000, anti-FANCD2 1/2000, anti-RAD51: 1/2000) and secondary antibody (Alexa 488-Anti-mouse IgG: 1/2000, Alexa 546-Anti-rabbit IgG: 1/1000) staining occurred consecutively at RT for 1 hour each. All antibodies were diluted in blocking buffer as described above. Nuclei were stained with 0.05µg/ml DAPI (Sigma-Aldrich) in PBS, for 15 min at RT. Cells were washed twice in 1x PBS with 0.1% Tween-20 after each step of staining: primary, secondary, and DAPI.

For non-S phase EdU incorporation assays, cells were treated with 10µM R0-3306(Sigma-Aldrich) 24hr prior to fixation and 10µM EdU was added 2hr prior to fixation. Staining was performed using the Click-iT Plus EdU Alexa Fluor 647 Imaging kit (Thermo Fisher Scientific), per manufacturer’s instructions prior to the commencement of additional immunofluorescent staining as outlined above.

All immunofluorescent images were captured using the Opera Phenix high-content imager with a 63X objective. Analysis was done using Harmony software (Revvity). Z stacks where maximum image projected and TIFs were called based on >50% overlap of telomere and DDR spots.

### Metaphase Spreads/FISH

Cells were treated with colcemid at 100 ng/ml for 1.5h, collected using trypsin and incubated at 37°C in prewarmed 75 mM KCl. The cells were spun down, and the KCl was removed. Cells were slowly and under agitation resuspended in a fixative of 3:1 methanol:acetic acid and stored overnight at 4°C. Next cells were spread, dropwise, onto cold, wet microscope slides and washed twice with 1 ml 3:1 methanol:acetic acid solution. Slides were then airdried and aged over night at room temperature. Slides were washed with PBS and then dehydrated in an ethanol series. Each slide received 100 µl of hybridization mixture, was denatured at 80 degrees for 5 min, and then hybridized overnight with Cy3 Tel-C probes (PNA Bio) and centromeric PNA probes (PNA Bio) at 4 degrees in a hybridization chamber. The next day, the slides were washed twice with 70% formamide, 10 mM Tris–HCl pH 7.2, and 0.1% BSA solution, then with twice with 0.1 M Tris–HCl pH 7.2, 0.15 M NaCl, and 0.08% Tween, and with DAPI (diluted 1:1,000 from 5 mg/ml stock) added to the second wash. Coverslips were mounted with ProLong® Gold Antifade Mountant (Thermo Fisher Scientific). All microscopy (IF and metaphase spreads) was imaged on a Nikon Eclipse TE2000-E epifluorescent microscope equipped with an Andor Zyla sCMOS camera.

### Alkaline Phosphatase Staining

After media was removed from hESCs grown on feeders, the plates were washed once in PBS and then treated with cold 4% PFA for 10min at RT. The cells were then washed twice with PBS. 100mM Tris-HCl (pH=9.5) was balanced in and left for 10 min at RT. One drop of each reagent from the Vector Red Alkaline Phosphatase Substrate Kit I (Vector Laboratories) was added to 6ml of 100mM Tris-HCl and added to the cells. Cells were developed at RT in the dark for ∼20 min until desired staining intensity was reached. Cells were washed once more and stored in PBS.

### High Throughput Imaging and Bayesian Regression Model

Stabilized POT1 mutant pools (14 days after editing) were single-cell dissociated and seeded onto 96-well plates using the Tecan Fluent liquid handler at a concentration of 30 cells/well based on an empirical 30% single-cell survival rate. Medium was changed every 3 days using Biotek EL406. At day 14, cells in each well were single-cell dissociated and split evenly into 3 parts using Tecan Fluent. Part 1 was replated onto new MEF plates for maintenance; part 2 was mixed with 2x lysis buffer to generate gDNA lysis and then NGS-genotyped as described above; part 3 was replated onto matrigel-coated plates to generate the 1st batch samples for imaging. The maintenance plates were then split the same way on day 21 to generate a 2nd batch of samples for imaging. Both imaging samples were fixed 3 days post plating, immunostained for TIFs and imaged as described above to measure the number of TIFs per cell. The TIF data from both imaging batches was combined and the frequency of TIF+ cells per well was calculated using TIF+ criteria >=10 TIF loci/cell. To reduce noise, wells that yielded <20 cells in imaging or failed in NGS genotyping were removed, and alleles that exhibited a frequency of <1% in all wells were binned into one low-frequency entry. The preprocessed allele frequency matrix and corresponding TIF+ frequency array were then used as input for a Bayesian regression model with a predefined assumption that the coefficients of all alleles follow a beta distribution (a=1, b=9) since the majority of alleles including all synonymous mutations don’t lead to TIF+. Scripts for allele classification and Bayesian regression are available on GitHub.

### Clonal isolation

Nucleofected cells were seeded onto 96-well plates at 10, 30 or 100 cells/well, alternatively mutant pools were seeded at 1, 3, 10 cells/well in hPSC medium with 10 µM Y27632. On day 12, plates were duplicated to generate gDNA lysis and then NGS genotyped as described above. Wells only showed one allele (>99% allele frequency) were expanded and genotype-confirmed to establish clonal cultures. To isolate frame-shift mutations, medium was supplemented with 1uM AZD6738 ATRi.

### Intragenic Synthetic Lethality

Each clone as well as the parental loHAP cells and diploid wild-type WIBR3 hESCs were nucleofected with a single sgRNA and accompanying ssDNA HDR oligopool as described above and cultured and sampled according to the deep scanning mutagenesis protocol. Allele classification was performed as previously described.

### ATM/ATR Inhibition

POT1 loHAP cells were nucleofected as described above, then immediately split into 3 biological replicates per 6 conditions with hPSC media supplemented as follows: Unsupplemented Control, Mock (0.1% DMSO), 1uM AZD0156 ATMi, 1uM AZD1390 ATMi, 10uM VE821 ATRi, or 1uM AZD6738 ATRi. Cells were cultured and sampled as described for 2 weeks before frame-shift analysis was performed.

### ATRi Withdrawal

Synonymous and frame-shift mutants maintained under ATRi were single-cell dissociated using accutase and seeded onto matrigel-coated 96-well plates at 4,000 cells/well with triplicates in hPSC medium +/- ATRi. Cells were fixed on day 5 using 4% PFA, stained with DAPI, and then imaged and counted on Celigo (Revvity).

### Telomere Length Detection

Genomic DNA was prepared as described previously.(*15*) Briefly, genomic DNA was digested with MboI, AluI, and RNase A overnight at 37°C. The resulting DNA was normalized, and 2 µg of DNA were run on 0.75% agarose (Seakem ME Agarose, Lonza), dried under vacuum for 2 h at 50°C, and denatured in 0.5 M NaOH, 1.5 M NaCl for 30 min, shaking at 25°C, neutralized with 1 M Tris, pH 6.0, and 2.5 M NaCl shaking at 25°C, 2× for 15 min. Then the gel was prehybridized in Church’s buffer (1% bovine serum albumin [BSA], 1 mM EDTA, 0.5 M NaPo4, pH 7.2, 7% SDS) for 1 h at 55°C before adding a 32P-end-labeled (C3TG2)3 telomeric probe. The gel was washed 3 × 30 min in 4× SSC at 50°C and 1 × 30 min in 4× SSC + 0.1% SDS at 25°C before exposing on a phosphor imager screen.

### Conditional CTC1 loop-out

Conditional CTC1 cell lines were created generated in WIBR3 hESCs (NIH stem cell registry 0079). Exon 5 deletion (Supplemental Figure 6A; Step 1) was generated via electroporation of CTC1 guide RNAs 1 and 2 cloned into a pX330 vector backbone. Cells were clonally isolated and genotyped to identify heterozygous CTC1^Δ/+^ clones. The repair template, consisting of a LoxP flanked exon 5 alongside an FRT flanked PGK-PURO cassette, was inserted (Step 2) by electroporation of CTC1 guide 3, which targets the junction on the Δ allele. After puromycin selection and clonal genotyping of successful CTC1^ΔPuro/+^ clones, the PGK-PURO cassette was removed (Step 3) via transfection with Flp mRNA to generate CTC1^F/+^ cells. Next, the remaining wild-type allele was removed (Step 4) by retargeting cells with CTC1 guide 1 and guide 2, generating CTC1^F/-^ cells. A CAGGS-ERT2-Cre-ERT2 expression cassette was then integrated at the AAVS1 locus in both CTC1^F/+^ and CTC1^F/-^ cells. To loop out the conditional CTC1 allele, cells were treated with 2µM 4-hydroxytamoxifen for 48hr.

### CTC1 Targeting in POT1 FS and Syn Cells

POT1 frame-shift and synonymous cells under ATRi were targeted with 2 sgRNA targeting exon 5 of CTC1. Nucleofection and clonal isolation was performed as described in the respective sections above. On day 19 post nucleofection, the plates duplicated on day 12 were fixed and stained with hNA and DAPI as outlined in the TIF Staining section. Cells were imaged and analyzed with a Celigo imager.

A bulk nucleofected population was kept in culture concurrent with the clonal plates and were then seeded at 1, 3, and 10 cells/well 14 days after nucleofection to clonally isolate cells at a later timepoint. These later timepoint clones were lysed and genotyped 26 days post nucleofection.

**fig. S1.**
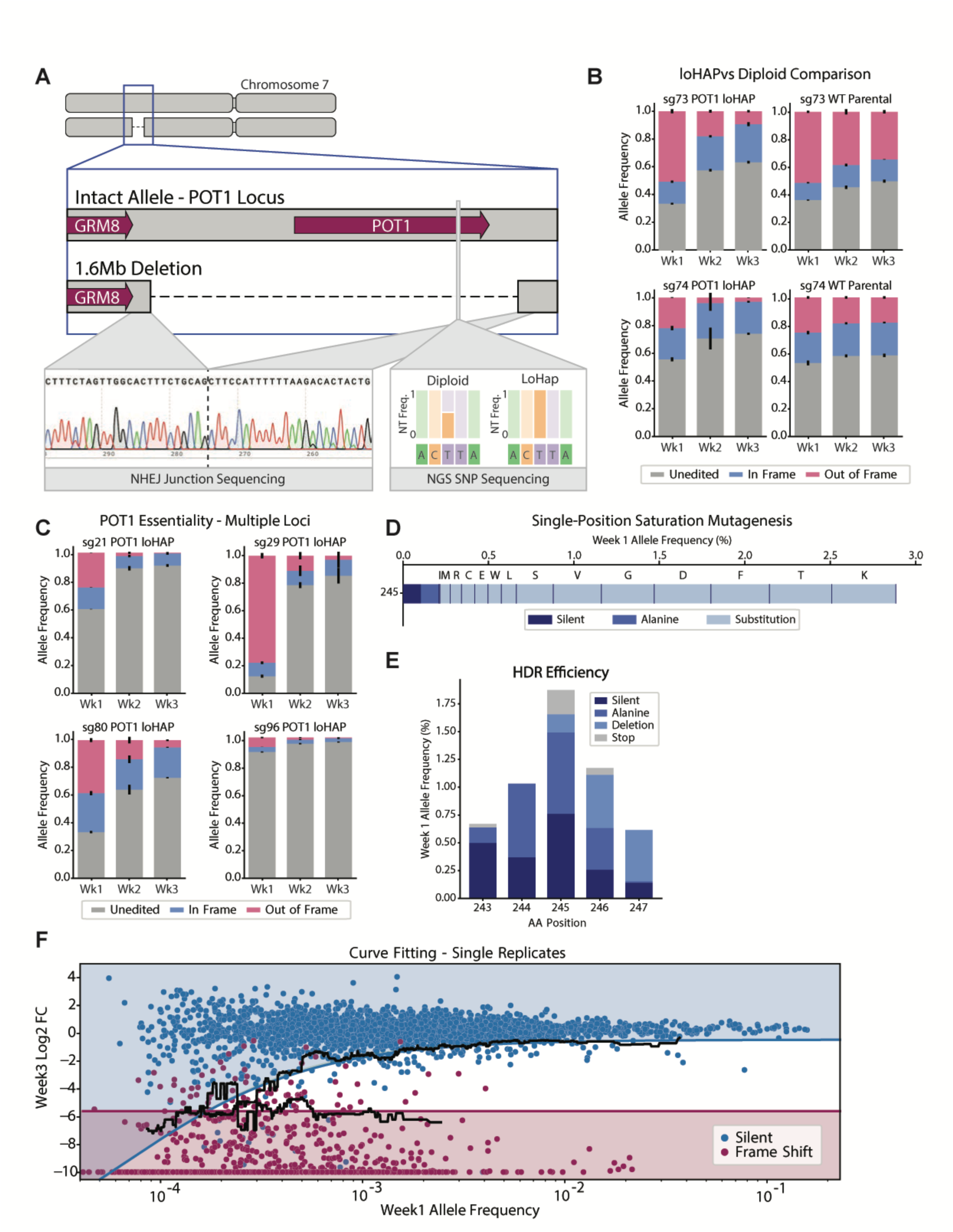
Development, characterization, and analysis strategies of deep scanning mutagenesis platform. (B) More detailed schematic of POT1 loHAP generation. Inset panels show sanger sequencing of the junction created by allele deletion (left) and next-generation sequencing of a single-nucleotide polymorphism located intronically between exon 15 and 16 (right). (**B**) Relative quantification of CRISPR/Cas9-mediated insertion and deletion alleles over three weeks in POT1 loHAP and diploid wild-type parental hESCs; three biological replicates, with error bars representing the standard error of the mean. Two unique sgRNAs shown. (**C**) Relative quantification of CRISPR/Cas9-mediated insertion and deletion alleles over three weeks in POT1 loHAP cells; three biological replicates, with error bars representing the standard error of the mean. Four unique sgRNAs shown. (**D**) Allele frequency of targeted mutations in a single-position saturation mutagenesis experiment using sgRNA 50. Substitutions other than alanine are indicated by single-letter amino acid abbreviations. 16/20 possible amino acid substitutions were detected with an overall targeted editing efficiency of ∼3% (**E**) Allele frequency of targeted mutations in a single-position mutagenesis experiment using sgRNA 50, establishing ∼6% allele incorporation for a 5 amino acid window surrounding the sgRNA cut site. (**F**) Scatterplot showing single biological replicates for silent mutations and frame-shift alleles generated during the screen. Black lines show the calculated 95th or 5th percentile (for frame-shift and silent mutations respectively) based on a scanning window of 180 data points. Curves were fit using the scipy optimize.curve_fit module. Allele Persistence/Depletion scores were calculated relative to these curves as described in the Materials and Methods section.

**fig. S2.**
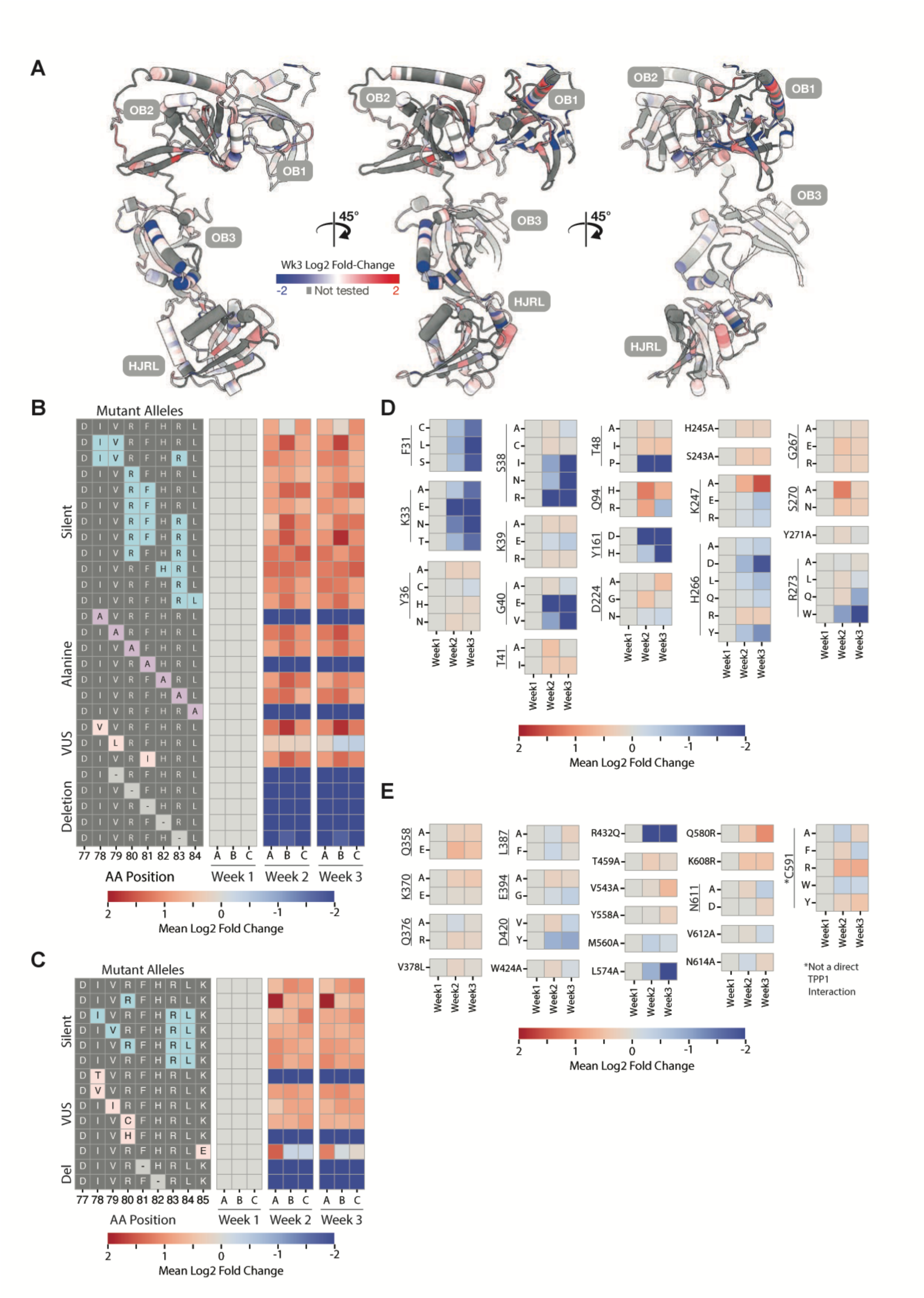
Mutations in POT1’s DNA-interacting or TPP1-interacting domains show varying levels of depletion. (**A**) POT1 AlphaFold2 (*60*) prediction colored by the week 3 log2 fold-change in allele frequency of alanine substitutions at the respective amino acid position. (**B-C**) Heatmaps showing log2 fold-change of three biological replicates across three weeks of sampling for POT-hole follow-up experiments. POT1 loHAP cells were targeted with a single sgRNA and an oligo pool designed for alanine/deletion scanning (B), or deeper VUS screening (C) and allele frequencies monitored for three weeks. (**D-E**) Heatmap showing mean log2 fold-change of three biological replicates across three weeks of sampling for residues which interact with telomeric DNA (*22*) (D) or residues which interact with TPP1 (*44*) (E) but were excluded from Figure 3 for having two or fewer unique missense mutations in our screen. Also included in 3B is C591 which does not directly interact with TPP1 but which abrogates POT1-TPP1 interactions when mutated (*33*). POT1-TPP1 contacts were identified by a Van der Walls overlap greater than or equal to −0.4 angstroms, ignoring interactions fewer than 4 bonds away.

**fig. S3.**
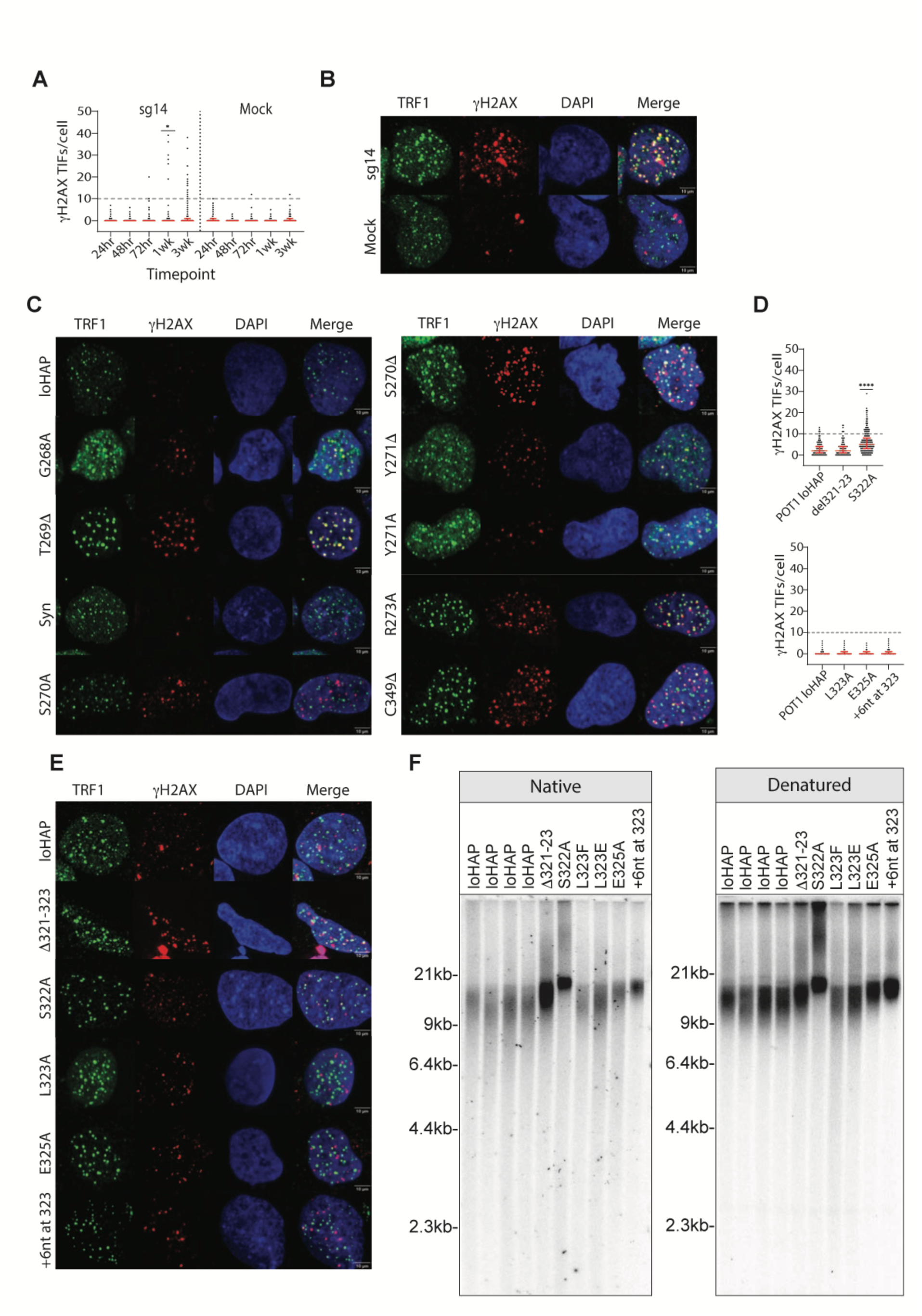
Disruption of POT1-CTC1 interactions results in telomere elongation and overhang extension. (A) Quantification of γH2AX TIFs over the course of 3 weeks for an additional sgRNA (sg14) targeting exon 7. Red error bars indicate median + interquartile range. Fisher’s exact test with FDR correction for multiple comparison of nuclei with =>10 TIFs was done between sg14 and Mock cells at each respective timepoint. * = p value <0.0332. (**B**) Representative images of γH2AX TIFs for sg14 alongside POT1 loHAP control cells at 3 weeks post editing. Scale bar, 10 μm. (**C**) Representative images of γH2AX TIFs in isolated mutant cells. Scale bar, 10 μm. (**D**) Quantification of γH2AX TIFs in different mutant cell lines. Two independent staining and imaging events were conducted and are plotted on separate graphs. Red error bars indicate median + interquartile range. Fisher’s exact test with FDR correction for multiple comparison of nuclei with =>10 TIFs was done for each cell line relative to POT1 LoHAP cells. **** = p value <0.0001. (**E**) Representative images of γH2AX TIFs in S322 mutants. Scale bar, 10 μm. (**F**) Telomere length analysis of clones with mutations around S322 under both native (left) and denaturing (right) conditions. Telomere probe: (CCCTAA)_3_

**fig. S4.**
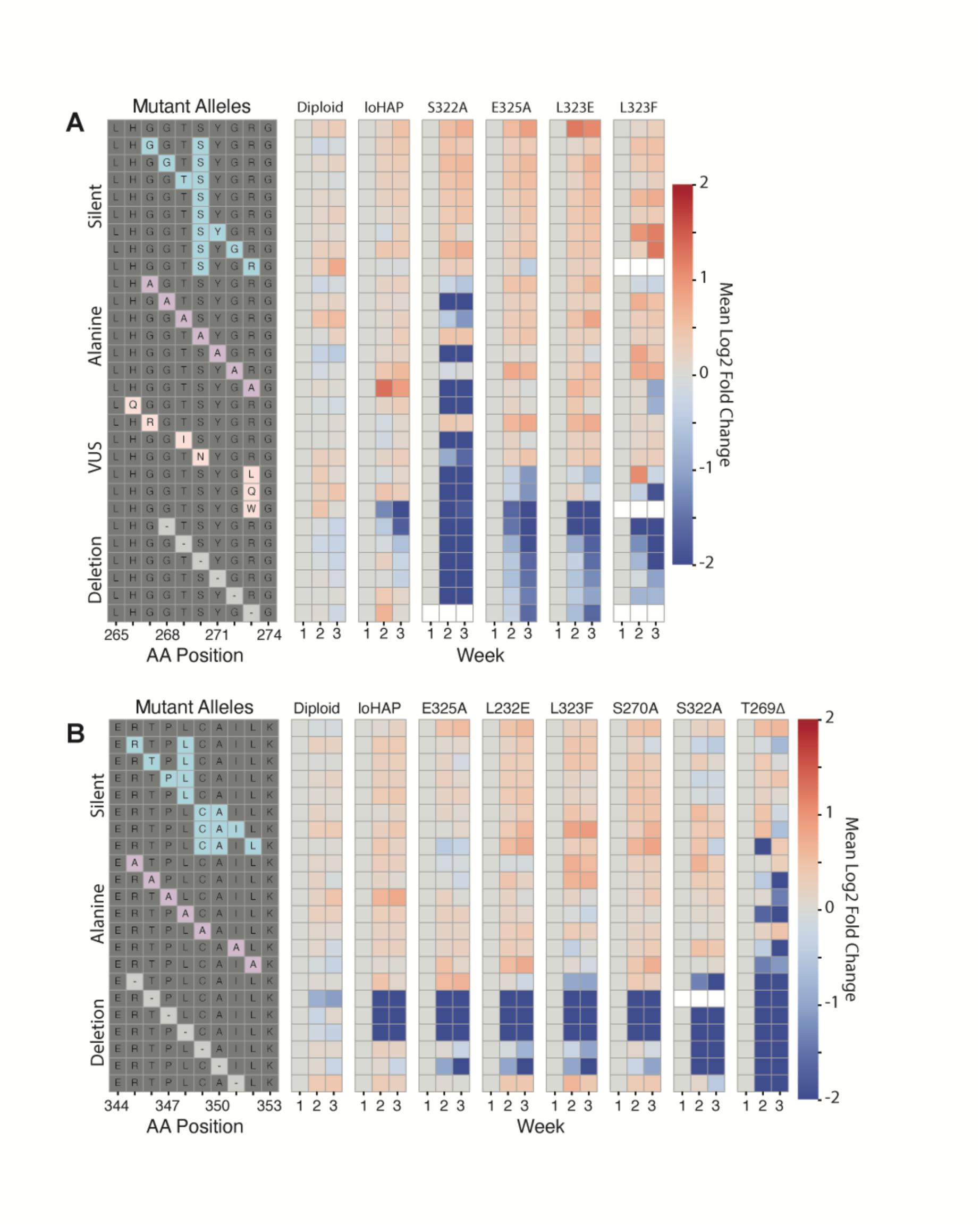
Intragenic synthetic lethality at multiple positions decreases cellular viability in TIF-positive backgrounds. Heatmap of intragenic synthetic lethality experiments using sgRNA 60 (**A**) and sgRNA 76 (**B**) showing mean log2 fold-change in allele frequency for three biological replicates across three weeks of sampling for each indicated cell line. Alleles introduced by the POT1 retargeting are grouped by category: silent, alanine substitution, or single amino acid deletion. Alleles which were not detected in all three biological replicates in an indicated cell line are shown in white.

**fig. S5.**
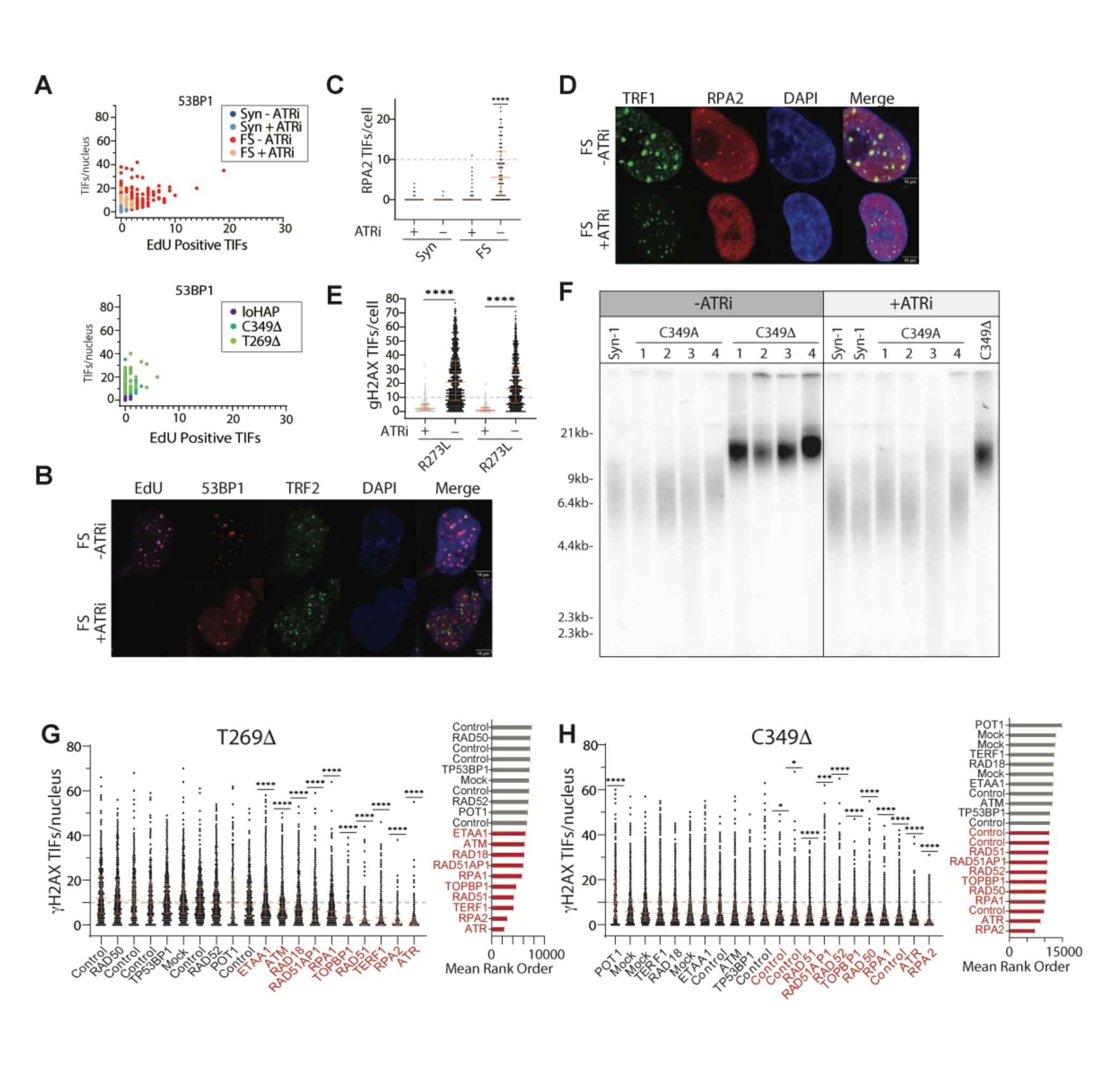
TIF phenotype in caPOT1 clones is ATR-mediated. (**A**) Frequency plots for sites of EdU positive 53BP1 TIFs (x-axis) vs. total 53BP1 TIFs per cell (y-axis). The top panel shows the frequency of distribution in loHAP, C349D and T269D POT1 mutant cells; the bottom panel shows frame-shift and synonymous mutations with and without ATRi. (**B**) Representative images for EdU and 53BP1 TIF colocalization in POT1 frame-shift cells ± ATRi. Scale bar, 10 μm (**C**) Quantification of RPA2 TIFs in frame-shift (FS) and synonymous (Syn) cells. Red error bars indicate median + interquartile range. (**D**) Representative images of RPA2 TIFs frame-shift (FS) and synonymous (Syn) cells. Scale bar, 10 μm (**E**) Quantification of γH2AX TIFs in two independent R273L mutant clones. Clones were not derived under ATRi but were placed under ATRi to show reduction of TIFs. Red error bars indicate median + interquartile range. Fisher’s exact test with FDR correction for multiple comparison of nuclei with =>10 TIFs was done between ATRi + and - cells. **** = p value <0.0001. (**F**) Telomere length analysis for C349A and C349Δ mutants derived under ATRi and without ATRi. Duplicate genotypes indicate independently derived clones. (**G**) Quantification of γH2AX TIFs in T269Δ cells at 7 days post targeted KO of 13 indicated genes. Red error bars indicate median + interquartile range. Fisher’s exact test with FDR correction for multiple comparison of nuclei with =>10 TIFs was done relative to mock transfected cells. ** =p value <0.0021, **** = p value <0.0001. Mean ranks were determined by assigning values for each nucleus depending on the number of TIFs present regardless of the experimental arm, and then determining the mean for each experimental arm based on the ranks assigned to each value in said arm. Mock nucleofected control cells are indicated as mock; non-nucleofected control cells are indicated as control. (**H**) As in G for C349Δ mutant cells

**fig. S6.**
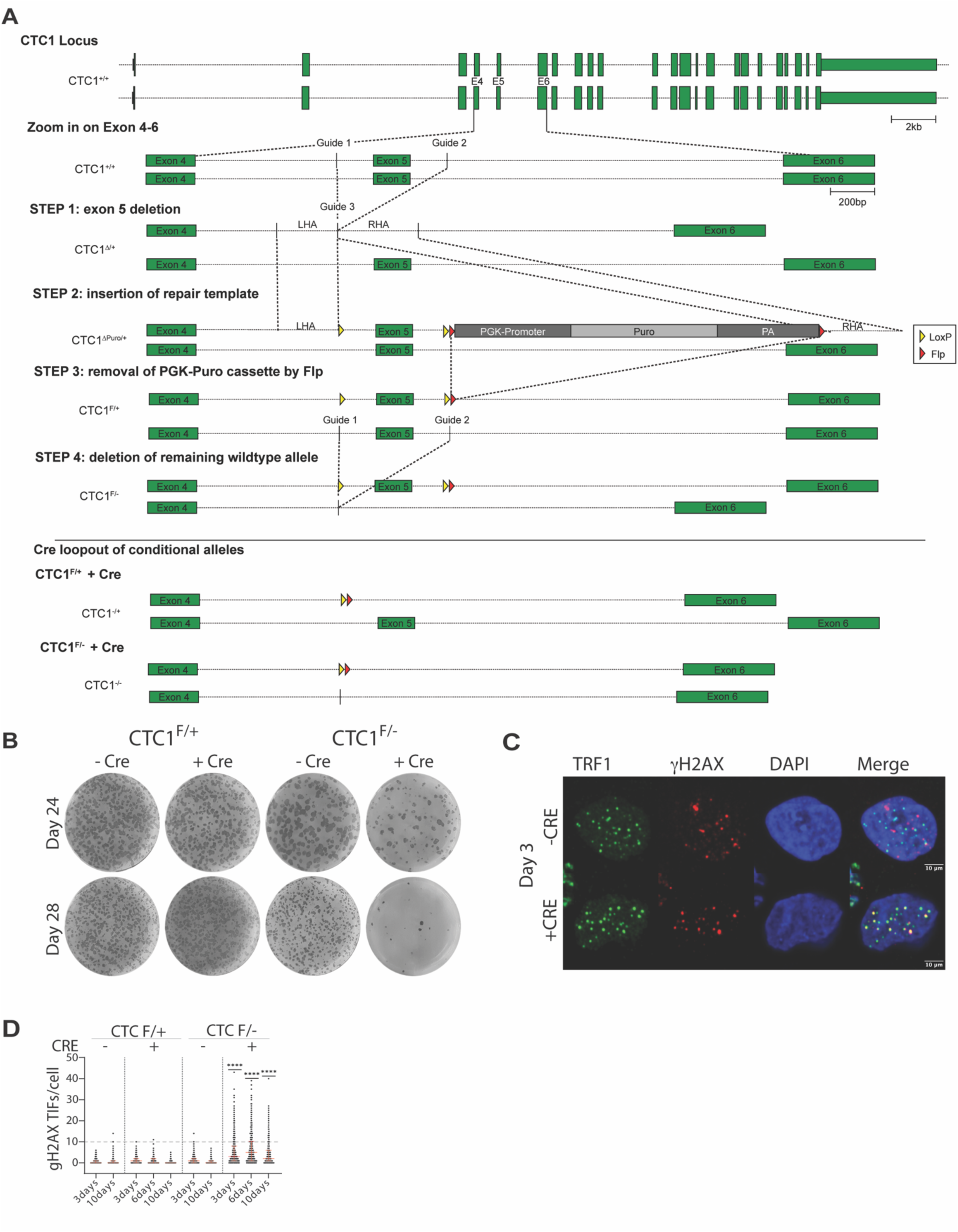
CTC1 knockout results in TIFs and gradual telomere shortening. (**A**) Schematic of genetic strategy used to engineer CTC1 conditional cells. (**B**) Alkaline phosphatase colony staining for conditional CTC1F/+ and CTC1F/- hESC cells after 24 and 28 days in culture with or without Cre induction. Cre was induced by treatment with 2µM 4-hydroxytamoxifen for 48 hours. (**C**) Representative images for CTC1F/- clones 3 days after addition of Cre. Scale bar, 10 μm. (**D**) TIF quantification for CTC1 conditional cell lines ± Cre. Red error bars indicate median + interquartile range. Fisher’s exact test with FDR correction for multiple comparison of nuclei with =>10 TIFs was done between corresponding CTC1 F/- and CTC1 F/+ cells. **** = p value <0.0001

**fig. S7.**
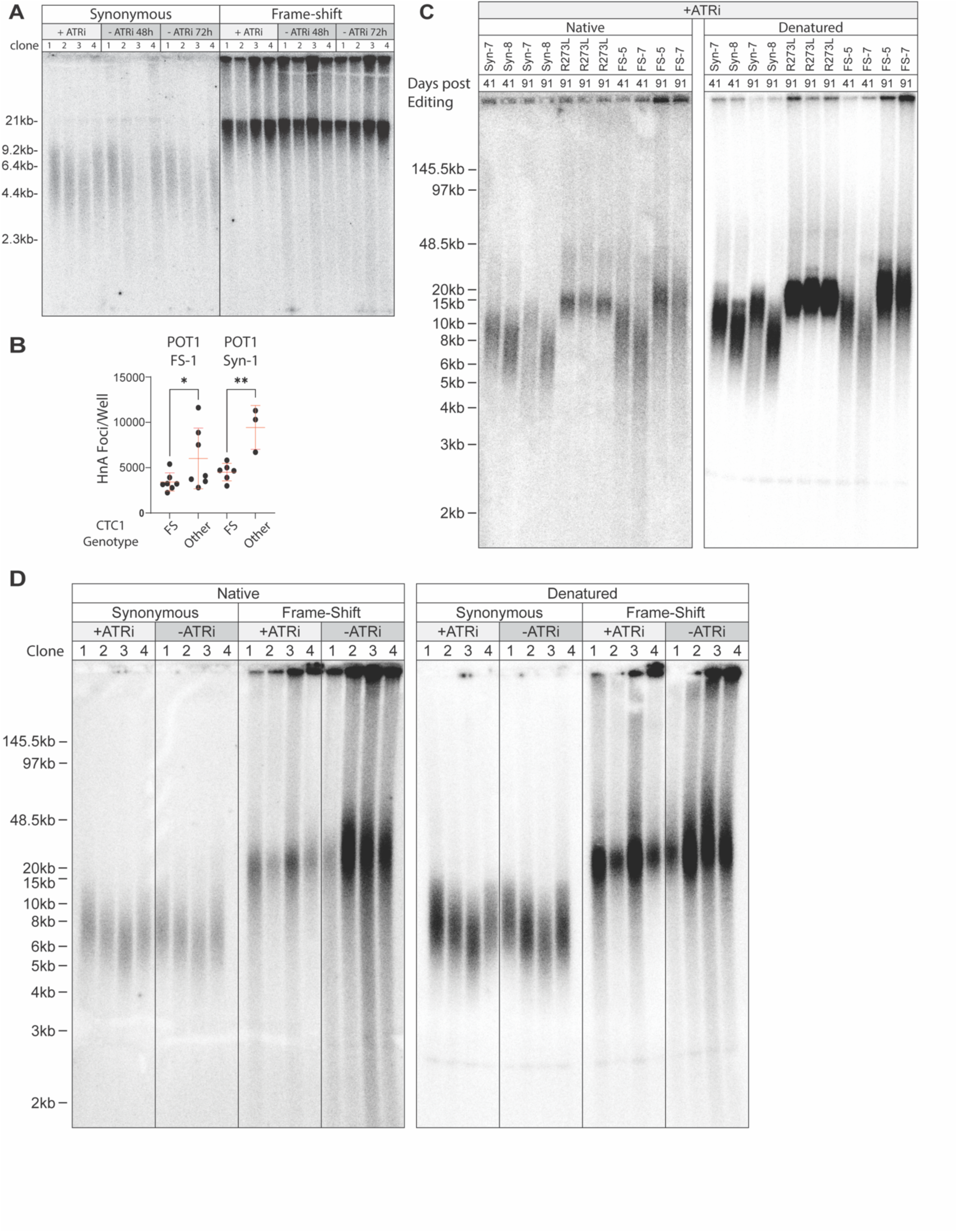
POT1 frame-shift clones show heterogeneous telomere elongation without overt overhang defects. (**A**) Telomere length analysis of four synonymous (Syn-1 through Syn-4) and four frame-shift (FS-1 through FS-4) clones four months post editing under the following conditions: ATRi, 48 hours after ATRi withdrawal, and 72 hours after ATRi withdrawal. (**B**) Quantification of human nuclear antigen foci per well for the “CTC1 FS” and “Other” clones generated in the POT1 FS-1 or Syn-1 background. “CTC1 FS” refers to clones which carry homozygous or compound heterozygous frame-shift mutations in CTC1. “Other” refers to any clone which carries at least one unedited or in-frame edited CTC1 allele. Red error bars indicate mean + SD. One-way ANOVA using Bonferroni’s multiple comparison test was used to indicate significance. * p value <=0.0332. (**C**) Telomere length and overhang analysis of synonymous (Syn), frame-shift (FS) and mutant clones derived under ATRi and sampled at indicated time points. Both native (left) and denaturing (right) conditions are shown. Duplicate R273L genotypes indicate multiple independently derived clones. Telomere probe: (CCCTAA)3 (**D**) Telomere length and overhang analysis of synonymous and frame-shift clones from panel A under ATRi and 72 hours post ATRi withdrawal. Both native (left) and denaturing (right) conditions are shown. Telomere probe: (CCCTAA)3

**Table S2.** Frame-Shift and Synonymous Clone Genotypes.

| **Exon Targeted** | **Genotype** | **Name in Figures** |
| --- | --- | --- |
| exon 11 | p.S270Ifs*19 |  |
| exon 11 | p.S270Ifs*19 |  |
| exon 14 | p.C349Vfs*4 | FS-1 |
| exon 8 | p.Y43Qfs*3 | FS-2 |
| exon 8 | p.Y43Vfs*21 | FS-3 |
| exon 8 | p.Y43Ifs*5 |  |
| exon 8 | p.Y43Lfs*7 | FS-4 |
| exon 10 | p.V229Mfs*6 |  |
| exon 10 | p.H226Mfs*6 |  |
| exon 11 | p.Y271* | FS-5 |
| exon 11 | p.Y271* | FS-6 |
| exon 11 | p.Y271* | FS-7 |
| exon 9 | p.Q160Gfs*5 |  |
| exon 9 | p.Q160Sfs*38 |  |
| exon 14 | p.C349,I351= | Syn-1 |
| exon 8 | p.Y43= | Syn-2 |
| exon 8 | p.Y43= | Syn-3 |
| exon 8 | p.Y43,S45= | Syn-4 |
| exon 11 | p.G272,R273= | Syn-5 |
| exon 11 | p.G272,R273= | Syn-6 |
| exon 11 | p.G272,R273= | Syn-7 |
| exon 11 | p.G272,R273= | Syn-8 |

**Table S3.**
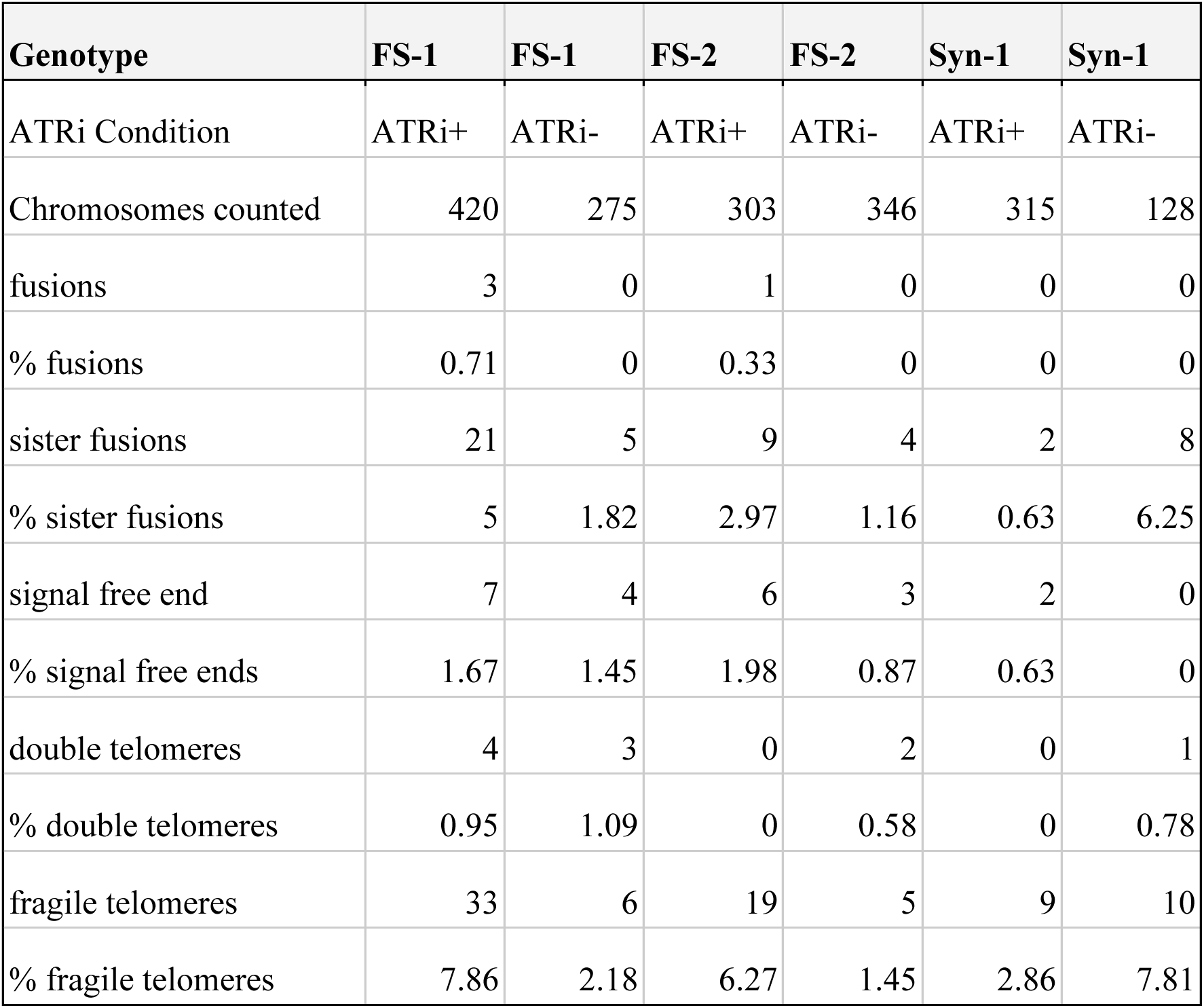
Quantification of Metaphase Spreads.

| **Genotype** | **FS-1** | **FS-1** | **FS-2** | **FS-2** | **Syn-1** | **Syn-1** |
| --- | --- | --- | --- | --- | --- | --- |
| ATRi Condition | ATRi+ | ATRi- | ATRi+ | ATRi- | ATRi+ | ATRi- |
| Chromosomes counted | 420 | 275 | 303 | 346 | 315 | 128 |
| fusions | 3 | 0 | 1 | 0 | 0 | 0 |
| % fusions | 0.71 | 0 | 0.33 | 0 | 0 | 0 |
| sister fusions | 21 | 5 | 9 | 4 | 2 | 8 |
| % sister fusions | 5 | 1.82 | 2.97 | 1.16 | 0.63 | 6.25 |
| signal free end | 7 | 4 | 6 | 3 | 2 | 0 |
| % signal free ends | 1.67 | 1.45 | 1.98 | 0.87 | 0.63 | 0 |
| double telomeres | 4 | 3 | 0 | 2 | 0 | 1 |
| % double telomeres | 0.95 | 1.09 | 0 | 0.58 | 0 | 0.78 |
| fragile telomeres | 33 | 6 | 19 | 5 | 9 | 10 |
| % fragile telomeres | 7.86 | 2.18 | 6.27 | 1.45 | 2.86 | 7.81 |

**Table S4.**
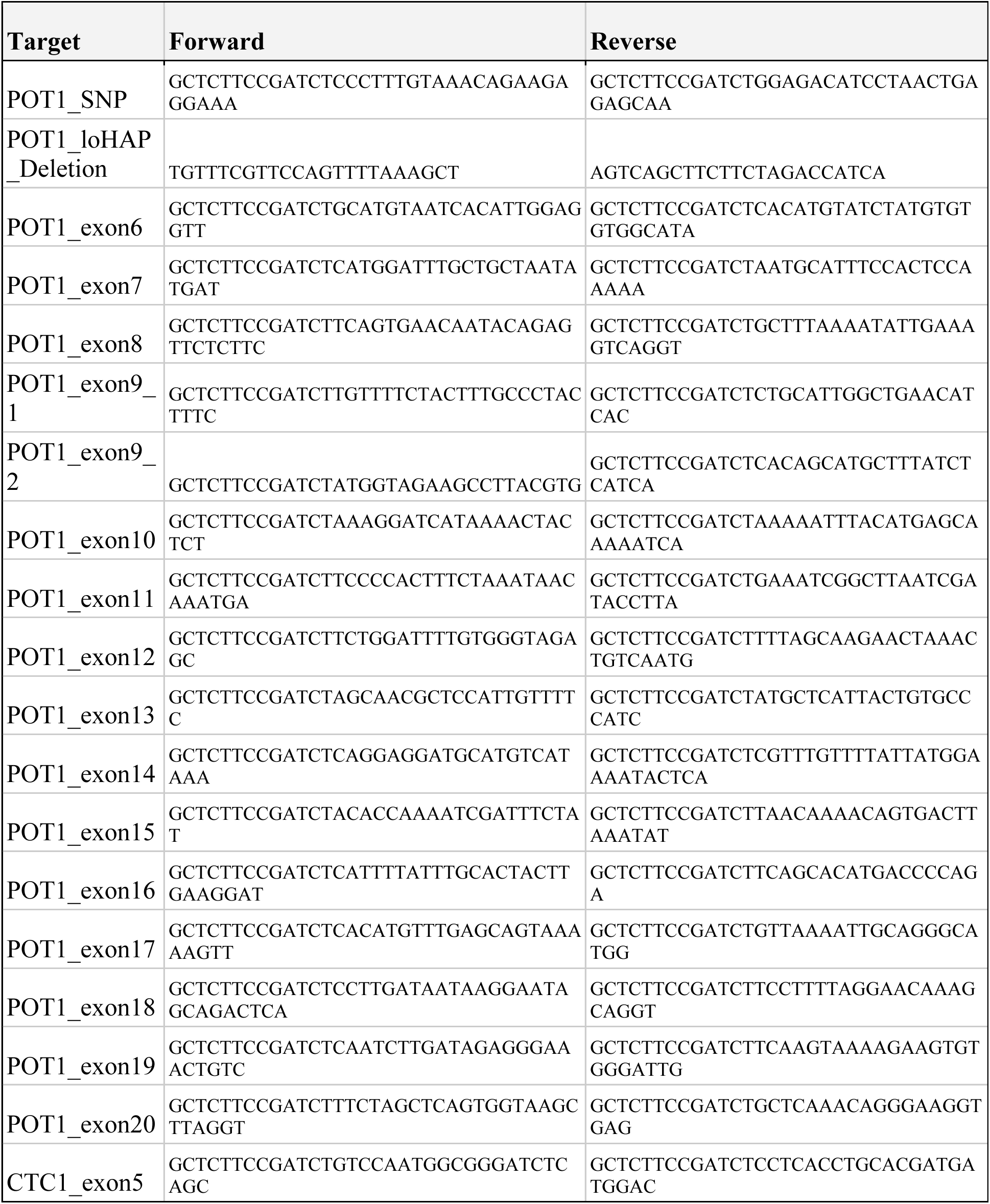
Primer sequences.

